# MixtureFinder: Estimating DNA mixture models for phylogenetic analyses

**DOI:** 10.1101/2024.03.20.586035

**Authors:** Huaiyan Ren, Thomas KF Wong, Bui Quang Minh, Robert Lanfear

## Abstract

In phylogenetic studies, both partitioned models and mixture models are used to account for heterogeneity in molecular evolution among the sites of DNA sequence alignments. Partitioned models require the user to specify the grouping of sites into subsets, and then assume that each subset of sites can be modelled by a single common process. Mixture models do not require users to pre-specify subsets of sites, and instead calculate the likelihood of every site under every model, while co-estimating the model weights and parameters. While much research has gone into the optimisation of partitioned models by merging user-specified subsets, there has been less attention paid to the optimisation of mixture models for DNA sequence alignments. In this study, we first ask whether a key assumption of partitioned models – that each user-specified subset can be modelled by a single common process – is supported by the data. Having shown that this is not the case, we then design, implement, test, and apply an algorithm, MixtureFinder, to select the optimum number of classes for a mixture model of Q matrices for the standard models of DNA sequence evolution. We show this algorithm performs well on simulated and empirical datasets and suggest that it may be useful for future empirical studies. MixtureFinder is available in IQ-TREE2, and a tutorial for using MixtureFinder can be found here: http://www.iqtree.org/doc/Complex-Models#mixture-models.

## Introduction

Differences in patterns of molecular evolution among sites in a sequence alignment have important consequences for the accuracy of phylogenetic analyses (Anderson and Swofford 2004; Blanquart and Lartillot 2008; Redmond and McLysaght 2021). This heterogeneity depends on many factors, including gene function, location on the chromosomes, codon position, the chemical properties of the encoded amino acids, protein structure, and many others (Kolaczkowski and Thornton 2008; Wang et al. 2008; Schrempf et al. 2020). A variety of methods have been proposed to account for this heterogeneity in phylogenetic analyses, including various approaches to modelling variation in rates of evolution (Yang 1994; Yang 1995; Soubrier et al. 2012; Kalyaanamoorthy et al. 2017) and variation in amino acid frequencies (Lartillot and Philippe 2004; Wang et al. 2018) among alignment sites. In this paper, we focus on approaches to modelling variation among sites in transition rates for DNA sequence alignments.

In Likelihood-based phylogenetic methods, transition rates are usually modelled by a single matrix, the Q matrix, which is computed from two parts – the exchangeability matrix (S) which specifies the instantaneous rates of changes between the four DNA bases (A, C, T, and G), and the frequency vector (F) which specifies the stationary frequencies of each of the four bases. Accounting for the heterogeneity among sites in a DNA sequence alignment typically involves solving two problems: determining how many Q matrices are required; and determining the type of Q matrices that should be used (e.g. by choosing from a list of Q matrices with different numbers of free parameters, such as the GTR model, the HKY model, and the JC model (Jukes and Cantor 1969; Hasegawa et al. 1985; Tavaré 1986)).

Partitioned models and mixture models have both been proposed to account for heterogeneity among sites for phylogenetic analyses of DNA sequence alignments (Gelman et al. 1995; Nylander et al. 2004). In a partitioned model, a sequence alignment is first partitioned into different data blocks (e.g., one block for each codon position in each protein-coding gene). Analyses are then performed assuming that the sites in each data block have evolved under a single substitution model (Nylander et al. 2004), while allowing different data blocks to have different models. Data blocks with sufficiently similar evolutionary models can be merged using a variety of information-theoretic approaches and algorithms, and the identity of each model can be selected at the same time (Lanfear et al. 2012; Lanfear et al. 2016). The upshot is the selection of a joint model in which the number and identity of Q matrices is determined, along with the assignment of each site in the alignment to a single Q matrix. Mixture models, like partitioned models, allow more than one substitution model to apply to an alignment. However, while a partitioned model requires each alignment site to be assigned to a specific model, a mixture model does not. Instead, a mixture model computes the likelihood of every site under every model, and optimises the relative weights of the models themselves (Le et al. 2008). The likelihood of each site in a mixture model is simply the weighted sum of site likelihoods for each model in the mixture (Pagel and Meade 2004). Simulations and empirical analyses have shown that both partitioned and mixture methods can improve the accuracy of phylogenetic analyses (Quang et al. 2008; Wang et al. 2008; Darriba and Posada 2015; Kainer and Lanfear 2015; Whelan and Halanych 2016; Crotty et al. 2018; Crotty et al. 2020; Redmond and McLysaght 2021).

Despite their popularity, partitioned models have an important limitation: their accuracy depends on both the accuracy and validity of our definitions of data blocks. Our ability to define meaningful data blocks is limited both by our knowledge of molecular evolutionary processes, and by the degree to which we can apply that knowledge to any given alignment. For example, it is common practice to partition alignments of protein-coding genes into data blocks by defining one data block for each codon position of each gene. This approach has repeatedly been shown to improve phylogenetic inference (Yang 1996; Brandley et al. 2005), compared to alternative ways of partitioning the data (e.g. by gene but not by codon position), but little is known about the validity of the assumption that a single model is adequate to describe the evolution of single codon position in a single gene. More generally, there is growing evidence that applying a single model to a single data block may limit the accuracy of our inferences (Wang et al. 2019; Crotty et al. 2020; Redmond and McLysaght 2021). In addition, many phylogenomic datasets consist of markers for which it remains unclear how to define meaningful data blocks. These include regions such as introns and intergenic regions, Anchored Hybrid Enrichment loci, Ultra Conserved Elements, and many others (Bejerano et al. 2004; Garcia-España et al. 2009; Lemmon et al. 2012). At the same time, it is clear that inaccurately defining data blocks, even when they exist, can mislead phylogenetic inference (Crotty and Holland 2022). Nevertheless, partitioned models are still more widely used than mixture models for DNA sequence alignments, likely because of the well-developed methods and software for estimating partitioning schemes and selecting models (Lanfear et al. 2012; Darriba and Posada 2015).

Mixture models offer an approach which circumvents many of the limitations of partitioned models, because they do not require users to pre-define data blocks. Mixture models are common in many phylogenetic settings. Variation in rates among sites is usually modelled as a mixture of different rate classes, e.g. where the rate of each class is estimated from a Gamma distribution (Yang 1994) or from a free-rate model (Yang 1995; Soubrier et al. 2012). Mixture models are also used to model variation in amino acid frequencies among sites in the CAT model (Lartillot and Philippe 2004; Quang et al. 2008; Minh et al. 2020b); variation in amino acid transition rate matrices among sites in models such as the EX3 and LG4X models (Le et al. 2008; Le et al. 2012); variation in branch lengths among sites in the GHOST model (Crotty et al. 2020); and variation among tree topologies in the MAST model (Wong et al. 2024). A recent study (Kapli et al. 2023) shows that for inferring deep phylogenies, DNA alignments are at least as useful as amino acid alignments. However, despite their potential utility mixture models are not commonly applied to the problem of modelling heterogeneity in transition rate matrices for DNA sequence alignments.

This study aims to evaluate the potential of mixture models of DNA Q matrices for phylogenetic inference. To this end, we first set out to test a key assumption made in most partitioned models – that a single Q matrix (i.e. a one-class model in the terminology of mixture models) is sufficient to model the evolution of a single data block. To do this, we compared the fit of a one-class model to that of a two-class mixture model on a large collection of the most commonly used data-blocks in phylogenetic analyses: individual codon positions from individual protein-coding genes. We then developed an algorithm which uses information-theoretic approaches to choose the optimal number of classes in a Q-matrix mixture model for any alignment. We call this algorithm MixtureFinder and implement it in IQ-TREE2 (Minh et al. 2020b). We evaluated the algorithm on simulated data and apply it to a range of small empirical datasets. We then applied it to a larger dataset of vertebrate loci, assembled to investigate the interrelationships of turtles, crocodiles, and birds (Chiari et al. 2012).

## New Approaches

MixtureFinder is an iterative algorithm implemented in IQ-TREE2 to estimate the optimal number of classes in a DNA mixture model (Figure 1). MixtureFinder starts by selecting the best single (non-mixture, i.e. one-class) substitution model (Q_1_) and rate heterogeneity across sites (RHAS) models (e.g. +G, +I+G, +R4, etc) using ModelFinder (Kalyaanamoorthy et al. 2017). MixtureFinder then iteratively adds classes to the existing mixture model, one at a time, while fixing RHAS models and tree topology. At the *k^th^* iteration (*k* ≥ 1), MixtureFinder tests all available DNA substitution models in IQ-TREE2 for the (*k*+1)^th^ class while fixing previously identified models for the 1^st^, 2^nd^, …, *k*^th^ classes and maximise the log-likelihood over all parameters in the Q-matrices in all classes, the parameters in the RHAS models, the tree branch lengths and class weights. The class weights 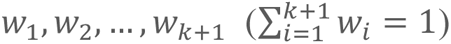 are optimised using the Broyden–Fletcher–Goldfarb–Shanno (BFGS) algorithm (Fletcher 2013). To help ensure model estimation accuracy, our algorithm employs a conservative and greedy strategy to initialise the (k+1)-class mixture model from the previously optimised parameters of the k-class mixture model (Supplementary Figure S1). This algorithm makes sure that the optimised likelihood of (k+1)-class model is at least as good as that of the k-class model. MixtureFinder then selects the best substitution model for the *(k+1)*^th^ class using the Akaike information criterion (AIC, Akaike 1974) or Bayesian Information Criterion (BIC, Schwarz 1978) score. Then, it uses the BIC (by default), AIC, or the likelihood ratio test (LRT, chi-squared test) to assess whether the (k+1)-class mixture model fits the data better than the previously determined k-class model. If it does, we accept the (k+1)-class mixture model as the better model, increase k by 1 and repeat this process. If it does not, we report k-class mixture model as the best model. Finally, MixtureFinder performs a full tree search given the mixture and RHAS model, optimises the model parameters and reports the model and the tree as the result.

**Figure 1.**
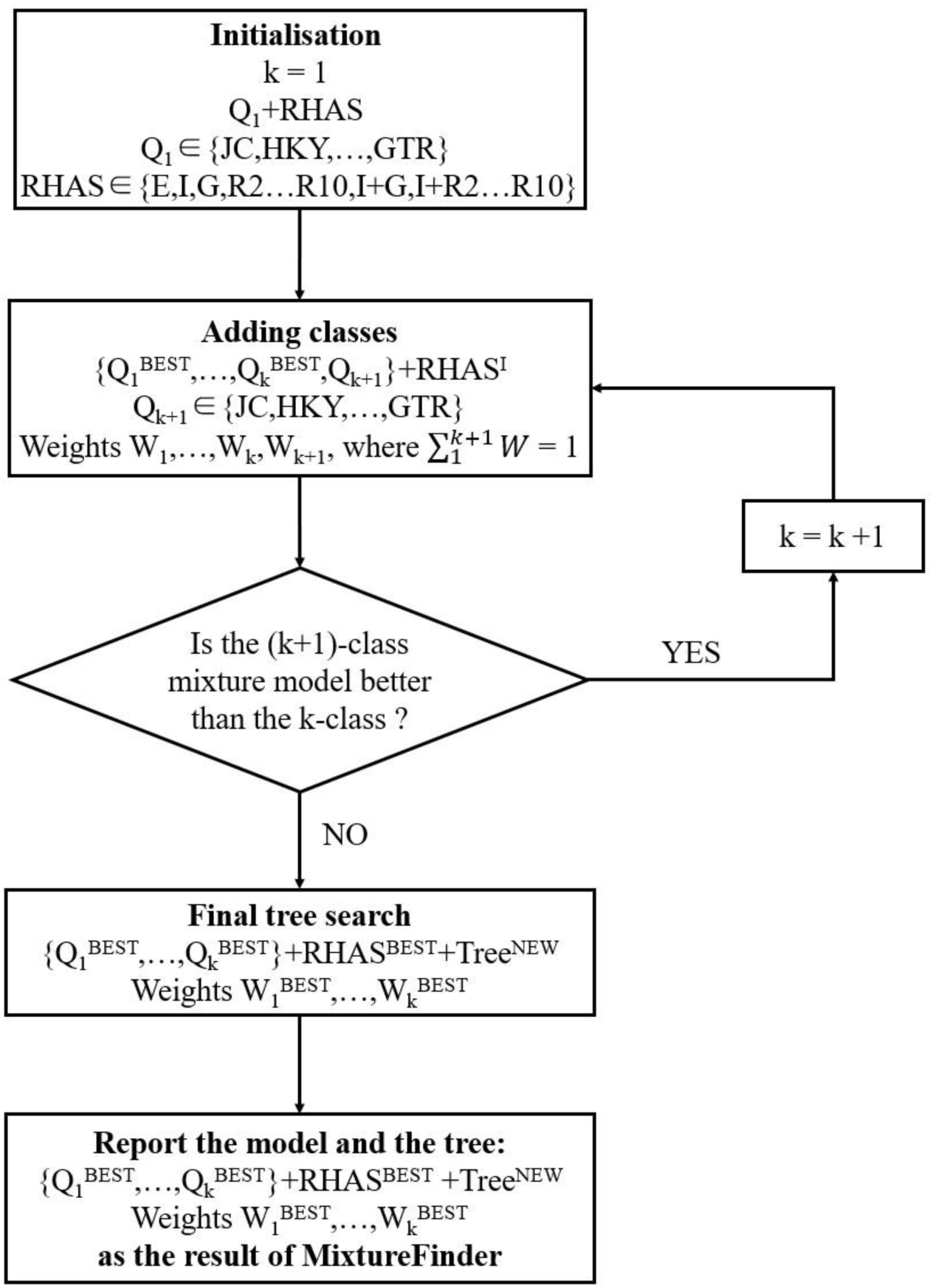
MixtureFinder algorithm design. MixtureFinder starts by fitting a 1-class model, adds classes one by one until the criterion do not support adding class. The optimal mixture model, RHAS model and the tree are reported as the output of the algorithm.

## Results

### Two-class mixture models almost always fit individual partitions better than one-class models

We first set out to test the assumption made in partitioned analyses that one-class substitution models (e.g., GTR+I+G) are sufficient to model the molecular evolution of a single partition (e.g., a single codon position from a single gene). To do so we simply ask whether the best two-class mixture model found by MixtureFinder is better than the best one-class model. We do this using both BIC and the LRT (p-value threshold 0.05) to determine the best mixture model. Our reasoning is that if a one-class model is sufficient to model the molecular evolution of a single partition, then partitioned models could, at least in principle, be sufficient to model the molecular evolution of any large collection of such partitions. On the other hand, if single partitions are often best-fit by a two-class mixture model over a one-class model, this would suggest that our ability to model molecular evolution via partitioning may be limited. We analysed three large DNA alignments of protein-coding genes from mammals, metazoans, and plants, splitting each into individual partitions based on genes and codon positions (Materials and Methods).

Across the 19,834 partitions from all three datasets, the vast majority were best fit by a two-class mixture model in preference to a one-class model (96.3% if the BIC is used for model selection, Figure 2A; 97.7% if the LRT is used for model selection, Supplementary Figure S2). The results differ somewhat across datasets and codon positions, with second codon positions having the smallest proportion of partitions best fit by a two-class model (89.8% if the BIC is used for model selection; 93.4% if the LRT is used for model selection), as expected if these partitions tend to evolve most slowly (and thus contain the least information) due to the constraints of the triplet genetic code.

**Figure 2.**
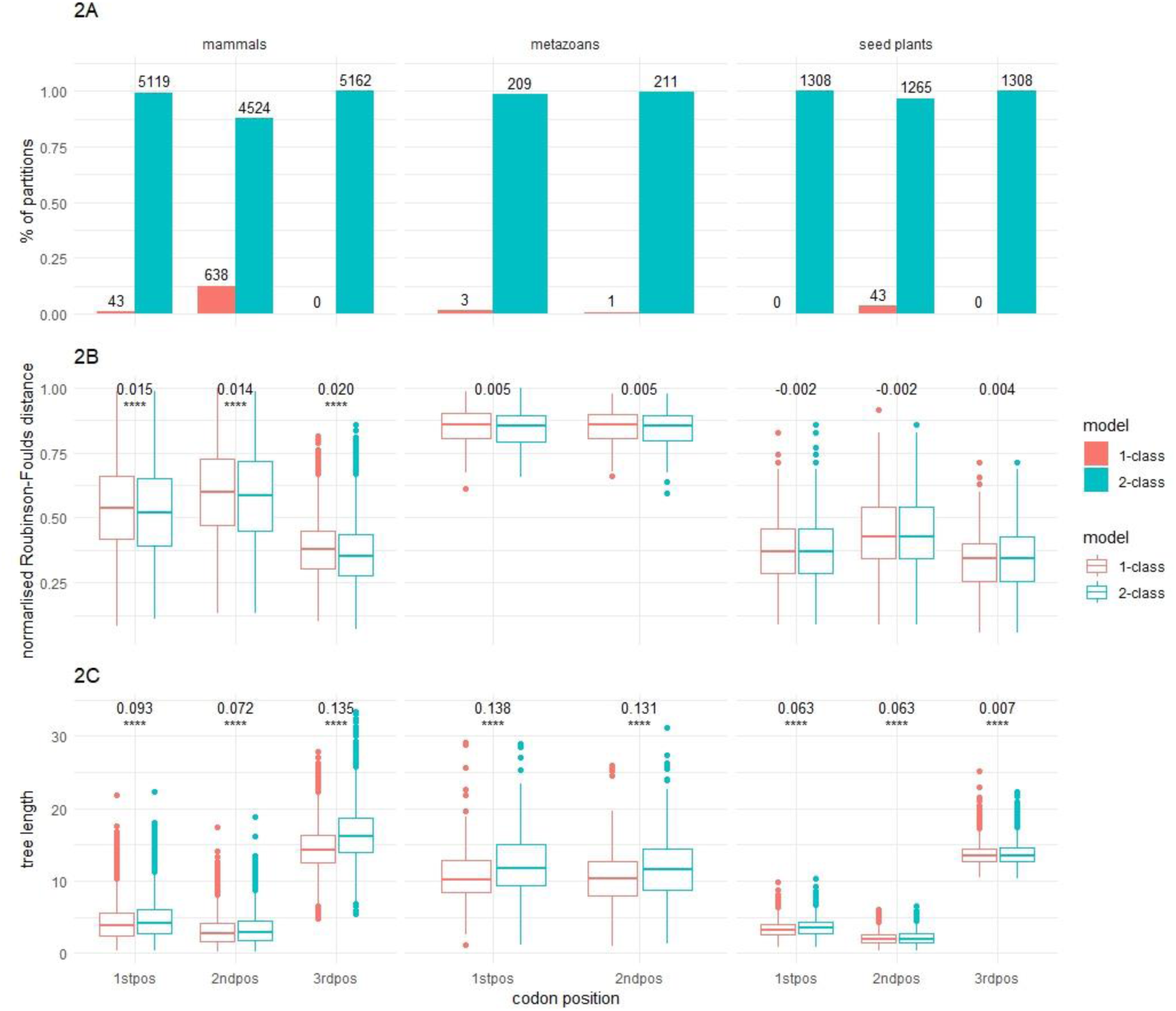
**A:** The proportion of partitions in which the 1- or 2-class model are better according to Bayesian Information Criterion (BIC). The exact number of partitions in which the 1- or 2-class model are better is shown on the top of each bar. **B:** The normalised Robinson-Foulds (nRF) distance between the commonly accepted tree and each of the 2-class tree and 1-class trees (for every partition). The difference of the average nRF (1-class trees – 2-class trees) is shown on the top of each pair of boxes. **C:** The tree length of the 2-class tree and 1-class tree (for every partition). The relative tree length difference of 2-class trees to 1-class trees (i.e. mean length of 2-class trees divided by mean length of 1-class trees) is shown on the top of each pair of boxes. The “*” marks represent the significance of the paired t-test between one-class and two-class models in 2B and 2C (0 ‘****’ 0.0001 ‘***’ 0.001 ‘**’ 0.01 ‘*’ 0.05).

### Single-partition tree topologies estimated from two-class mixture models are slightly closer to the estimated species tree than those estimated from one-class models

We next sought to understand the impact of mixture models on trees inferred from individual partitions. To do this, we selected every partition that was best-fit by a two-class model and estimated two maximum likelihood trees: one using the best two-class model, and another using the best one-class model estimated for that partition. We compared the trees for each partition in two ways. First, we compared the topologies of the trees estimated under the one- and two-class models by calculating normalized Robinson-Foulds (nRF) distance (Robinson and Foulds 1981) between each tree and the species tree of the same taxa estimated from the entire concatenated dataset (Materials and Methods). We reasoned that despite the limited information contained in each partition (and thus the high stochastic error of any tree estimated from one partition), if model fit is indicative of tree accuracy, then the tree estimated from the best-fit model (the two-class model in this case) may be closer to the estimated species tree than those estimated from the one-class model. We acknowledge, of course, that this does not account for biological processes that may cause partition-trees to differ from the species tree, such as hybridisation or incomplete lineage sorting. Second, we compared the total length of trees estimated from the one- and two-class models.

Figure two shows that where the two-class model is the best fit, two-class mixture models tend to result in trees which are slightly closer (Figure 2B) to the reference tree and slightly longer (Figure 2C) than those derived from one-class models. As before, these results differ somewhat among datasets and codon positions. For example, two-class trees are on average 0.02 nRF units closer to the concatenated tree for 3^rd^ codon positions in mammals (corresponding to the two-class tree being on average 3.6 splits more similar to the 90-taxa reference tree). On the other hand, two-class trees are on average 0.004 nRF units closer to the concatenated tree for 3^rd^ codon position in seed plants. Paired t-tests (two-tailed) were conducted and showed that two-class model results in the tree topology significantly closer to the reference tree, for all codon positions in the mammals. Scatter plots of the same data (Supplementary Figure S3) reveal few outliers, suggesting that these small differences are quite consistent among different partitions from the same dataset. Similarly, the 3^rd^ codon position trees are on average 1.95 and 0.09 substitutions per site longer for the two-class than the one-class trees for mammals and seed plants respectively, but this difference is smaller for the 1^st^ and 2^nd^ codon positions. The paired t-tests showed that two-class model results in a significantly longer tree for all these cases.

### MixtureFinder successfully recovers mixture models on simulated data with sufficient information

To assess the accuracy of MixtureFinder, we tested it on a range of simulated alignments under two different simulation schemes (Materials and Methods) and applied MixtureFinder to each simulated alignment in two ways, in both cases using the BIC as the model selection criterion) In the first application of MixtureFinder, which we call Q-mixture below, we test all available DNA sequence models in IQ-TREE2 (JC, HKY, GTR, etc.) when adding each mixture class. In the second application of MixtureFinder, which we call GTR-mixture below, we simply use the GTR model for each mixture class. We reasoned that the latter method may be a lot faster, and may sacrifice little or no accuracy, since if an alignment is best fit by a mixture of Q-matrices then perhaps it is unnecessary to examine substitution models which are more constrained than the GTR model (Abadi et al. 2019; Guimarães Fabreti and Höhna 2023).

Assessing the runtime and memory requirements of the GTR-mixture revealed that the GTR mixture is always faster than the Q-mixture (with the difference increasing with the number of estimated classes, up to an almost 6-fold speedup on the most complex simulations with 10,000 sites and five simulated model classes, which takes on average 3.73 hours to estimate the GTR-mixture but 22.02 hours to estimate the Q-mixture), and marginally more memory efficient (Supplementary Figure S7). This is because the GTR-mixture only requires a single model to be evaluated at each step (Supplementary Figure S1).

We compared the estimated models and trees to those used to simulate the alignment in four ways: the number of classes, the model parameters (using the cumulative density functions, Materials and Methods), the tree topology (using the nRF distance between the estimated and simulated trees), and the tree length. For clarity, we only show the results of the shortest and longest simulated alignments here (Figures 3-5), while figures including all simulated alignment lengths are shown in the Supplementary Information (Figures S4, S5 and S6). The ability of MixtureFinder to recover the number of model classes used to simulate an alignment depends on the amount of information in that alignment (Figure 3). Here, each alignment has 100 sequences. For example, in the simulation scheme in which the models in each class differ substantially (scheme 1), MixtureFinder usually recovers the number of model classes used to simulate the data even with the shortest alignment lengths we simulated (1000 sites; Figure 3). But in the simulation scheme in which the models in each class are more similar (scheme 2), MixtureFinder frequently underestimates the number of model classes used to simulate the data. Similarly, the performance of MixtureFinder improves with longer alignments (Figure 3). The differences between using the Q-mixture and GTR-mixture approaches are small regardless of the simulation settings (Figure 3).

**Figure 3.**
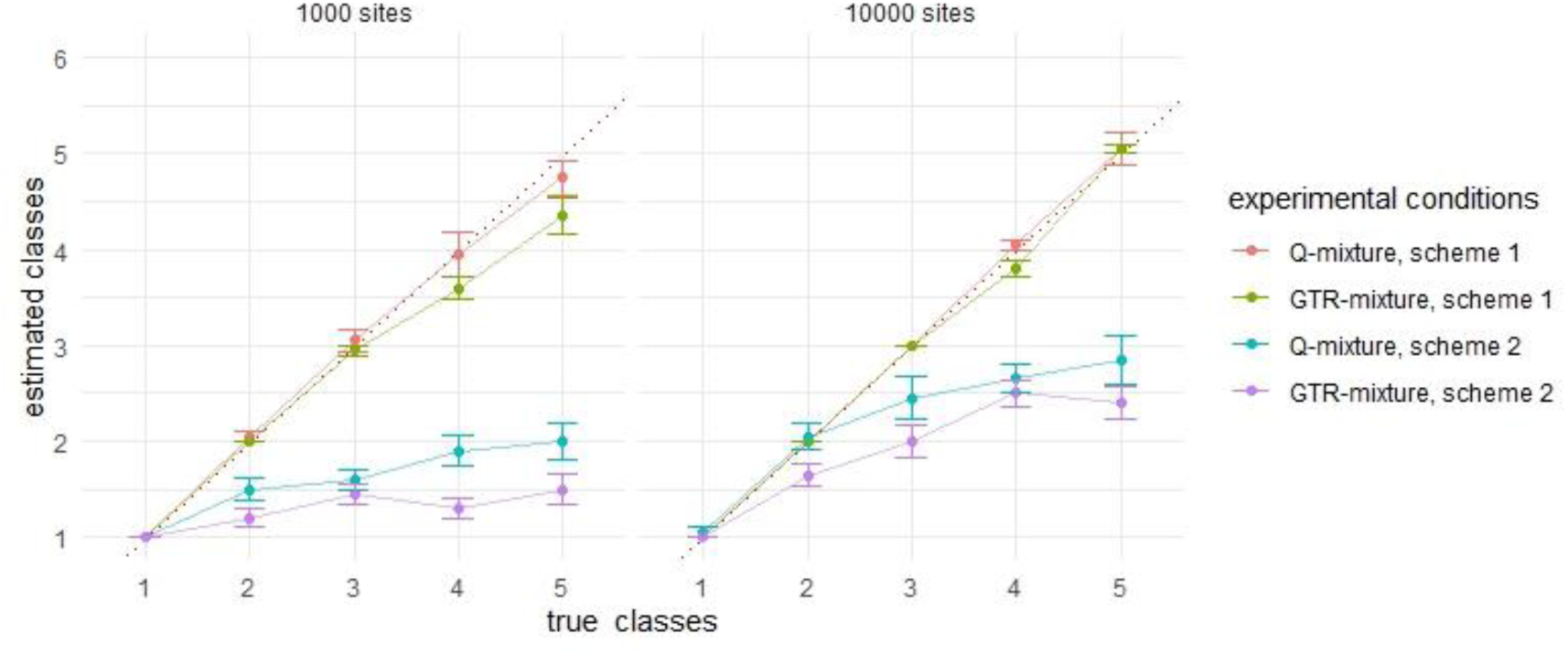
Estimation of the number of model classes. The red dashed line (y = x) represents that the estimated number of classes equals to the true number. The mean value and the corresponding standard error of the mean (error bar) of each experimental condition are shown with different curves.

The integrated squared error (ISE, Baños et al. 2024) between the parameters of the simulated models and estimated models shows that both the exchangeability matrix and the frequency vectors are recovered with high accuracy using MixtureFinder, regardless of the simulation conditions (Figure 4). As with the number of model classes, MixtureFinder produces more accurate results with longer alignments (e.g. higher error for 1000 bp alignments than for 10,000 bp alignments in Figure 4) and when the simulated models are more different from each other (e.g. higher error for simulation scheme 2 than for simulation scheme 1 in Figure 4). The ISE remains stable regardless of the number of simulated classes in the mixture models. An one-tailed paired t-test showed that for some simulation conditions the Q-mixture approach is significantly more accurate (i.e. recovers parameters significantly lower errors) than the GTR-mixture approach (Figure 4). This happens more frequently with shorter alignments (1000 bp). The most extreme case is observed when data were simulated under a single-class within simulation scheme 2, where some Q matrices have equal base frequencies that are estimated inaccurately with GTR models, compared to the Q-mixture approach which often selects models (such as the JC model) where base frequencies are identical. The ISE of mixture models is always better than the non-mixture (one-class) models when the true models are mixture models, no matter how many classes were simulated in the mixture models (Figure 4). In simulation scheme 1, the ISE of mixture models is around ten times lower than non-mixture models. In simulation 2, the difference is smaller, especially for 1000 bp alignments (around 1.5 times lower on average).

**Figure 4.**
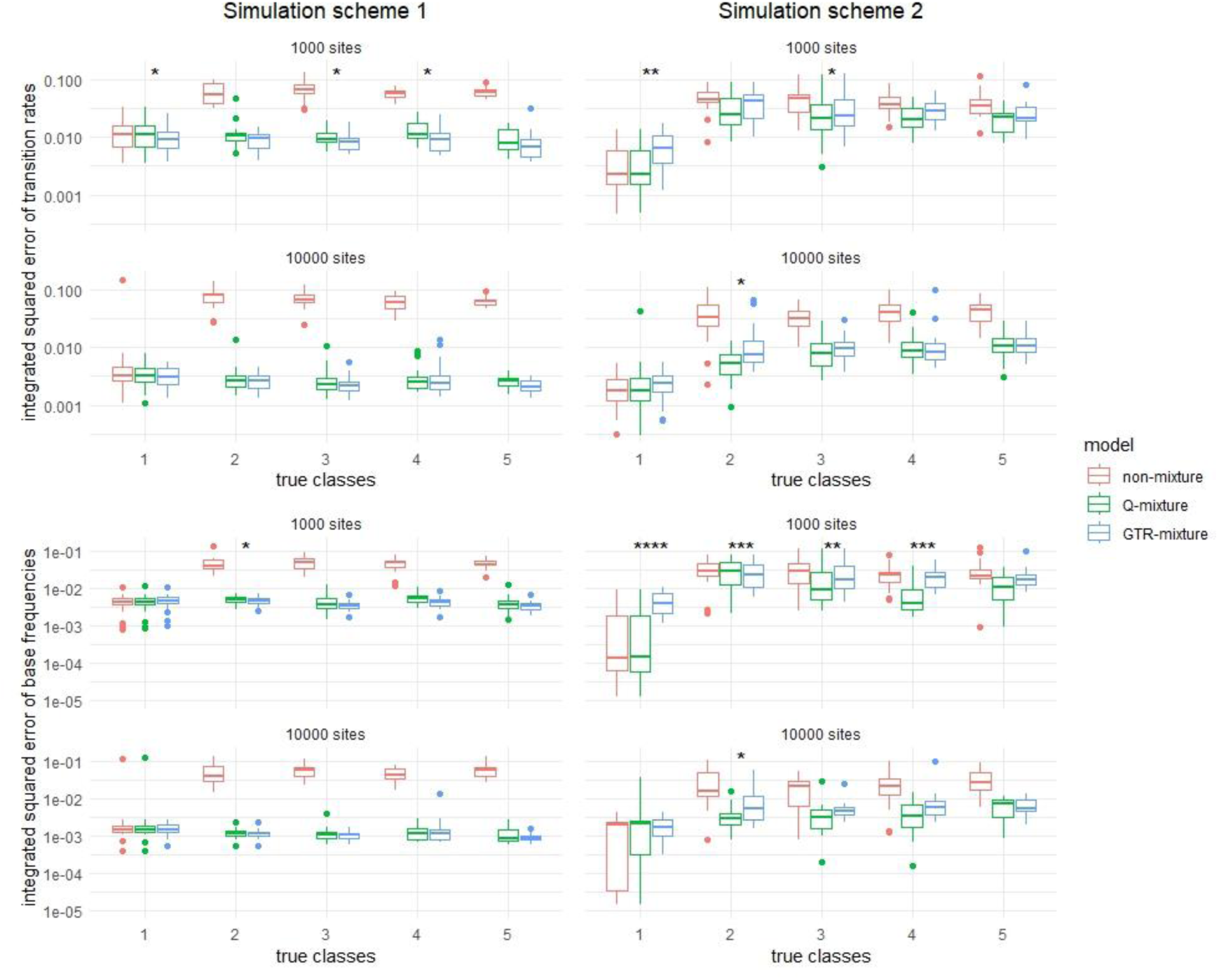
Integrated Squared Error (ISE) between parameters in estimated model and simulated model. The mixture models (green and blue bars) have smaller ISEs to the simulation than the non-mixture models (red bars). The “*” marks represent the significance of the paired t-test between Q-mixture and GTR-mixture models (0 ‘****’ 0.0001 ‘***’ 0.001 ‘**’ 0.01 ‘*’ 0.05).

Trees estimated by models selected with MixtureFinder have similar topologies to those estimated using one-class models, but much more accurate tree lengths (Figure 5). The nRF distances show that the topologies of all inferred trees are similar for any given combination of simulation settings. In contrast when data are simulated under a mixture model, fitting a one-class model tends to lead to underestimation of the simulated tree length. This is sometimes substantial for simulation scheme 1 where tree lengths were underestimated by up to 15% by the one-class models. This difference was much smaller for simulation scheme 2, where the underestimation is around 2%.

**Figure 5.**
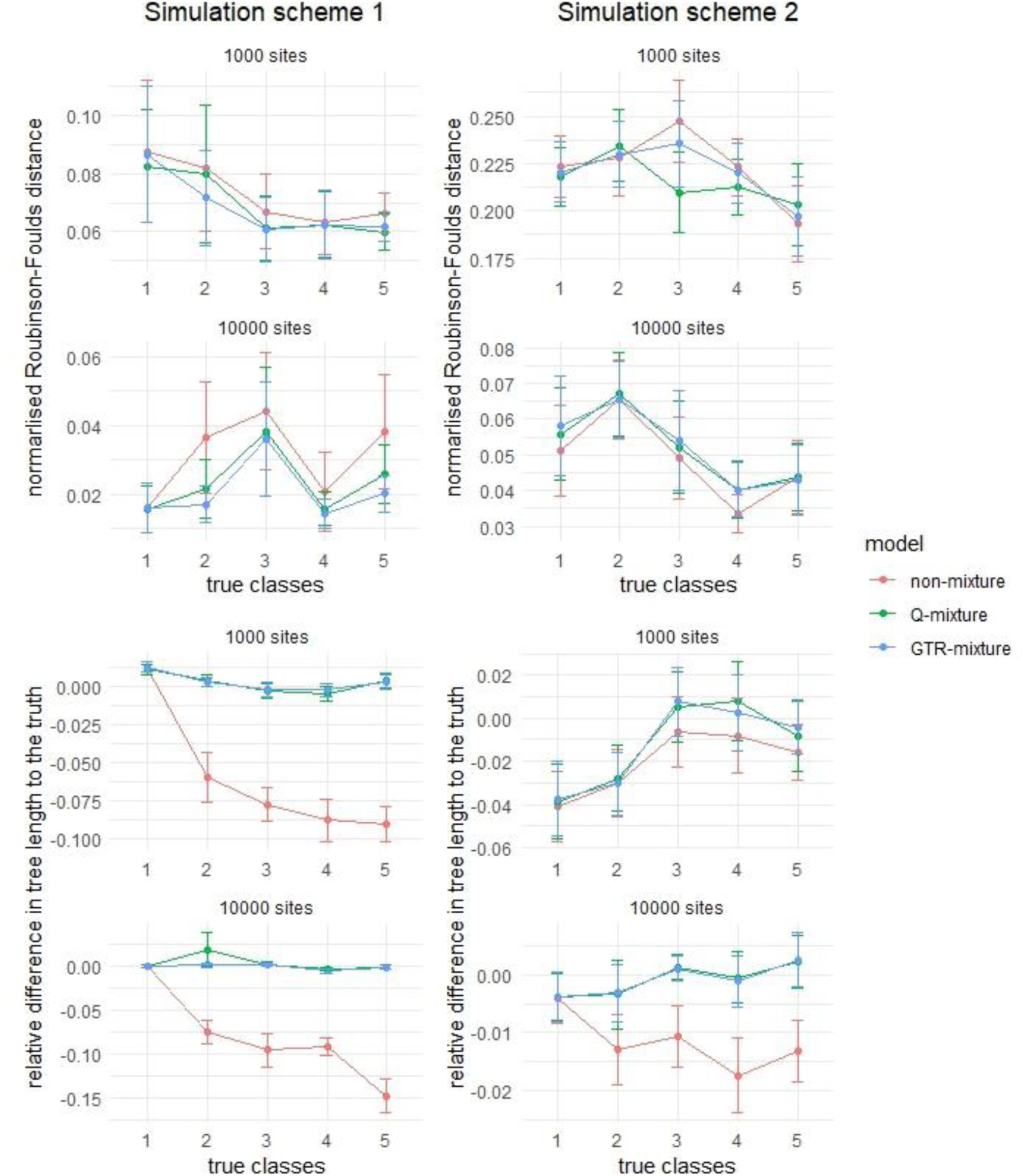
The normalized Robinson-Foulds (nRF) distance and relative difference in tree length between estimated model trees and true trees. The mean value and the corresponding standard error of the mean (error bar) of each experimental condition are shown with different lines. The nRF of mixture model trees to the simulated trees are slightly smaller than the non-mixture model in only simulation scheme 1, 10000-site alignments. In both simulation schemes, the mixture models recover the tree lengths that are closer to the lengths of the simulated trees, while the non-mixture models underestimate the tree lengths.

Finally, we assessed the ability of mixture model optimisation to recover simulated class weights when the true number of classes is known, by directly estimating the N-class GTR-mixture model, where N is the number of simulated number of classes (Supplementary Text 2). We matched the estimated and simulated mixture model (Supplementary Figure S8) and calculated the root mean squared error (RMSE) of class weights. The results showed that the average RMSE of weights was less than 0.1 in simulation scheme 1. Meanwhile, the RMSE in simulation scheme 2 ranged between 0.1 and 0.3, indicating poorer performance compared to scheme 1 (Supplementary Figure S8). This can be explained by the higher difficulty of the scheme 2 (Materials and Methods).

### The Likelihood Ratio Test gives different results to using a Likelihood Ratio Statistic (LRS) threshold

One limitation of the Likelihood Ratio Test (LRT) is its reliance on a chi-squared distribution, which is known not to hold for mixture hypotheses (Mitchell et al. 2019). Because of this, we compared the LRT to using a simple threshold for the Likelihood Ratio Statistic (LRS), when data were simulated under the null model. We ran MixtureFinder up to two-class models on 1000 datasets simulated under one-class models (see Materials and Methods). We plotted the distribution of the Likelihood Ratio Statistics (LRS) in 1000 datasets for the comparison between the two-class and one-class models on each dataset, highlighting the 95^th^ percentile of LRS and the 0.05-level chi-squared test threshold with the degree of freedom that corresponds to each model (Supplementary Figure S9). The results included six different Q matrices in the second mixture class: JC, K2P, TPM3, HKY, TIM and GTR. Consequently, the two-class models (alternative models) have 1, 2, 3, 5, 7 and 9 more degrees of freedom than the one-class models (null models) respectively (including the parameter for the additional class weight). With JC in the second class, the 95^th^ percentile of LRS was 3.55, close to the 0.05-level of chi-squared test threshold with 1 degree of freedom (3.84), and similar results were observed for K2P and TPM3. As the degrees of freedom increased, the LRS percentile became larger than the chi-squared test threshold. With GTR in the second class, the 95^th^ percentile of LRS is 19.67, larger than the chi-squared test threshold (16.92). The proximity of these values suggests that, with an LRT p-value threshold of 0.05, more than 5% of one-class models might be rejected in these 1,000 one-class simulations. In this experiment, 986 of 1000 samples have JC as the optimal (selected by BIC, as we explained in “New approaches”) Q matrix in the second mixture class, and the other 12 datasets have K2P or TPM3 in the second class. Overall, 0.57% of one-class models are rejected the 1000 samples.

### Mixture models fit empirical datasets better than one class models, and change phylogenetic conclusions

We applied MixtureFinder to a dataset that contains 16 vertebrate taxa with 248 genes and around 186 thousand base pairs and focuses on the relationships among turtles, crocodiles, and birds. (Chiari et al. 2012). The original study demonstrated that the topology which relates birds, crocodiles, and turtles can depend on the model of evolution used, with an un-partitioned one-class model grouping crocodiles and turtles, but partitioned models grouping crocodiles and birds. To better understand the effects of mixture models on empirical phylogenetic inference, we not only ran MixtureFinder to generate the tree under the model with the optimal number of classes, but also estimated a tree under a mixture model with every number of classes up to the optimal number, including the one-class (i.e. non-mixture) model. The optimal mixture model determined by MixtureFinder on the vertebrate dataset had seven classes (Figure 6). The seven-class model has a BIC score of 2091558, compared to a score of 2191133 for the initial one-class model. Trees inferred under the one- and two-class models group together crocodiles and turtles with bootstraps of 100 and 55 respectively (Figure 6B). However, trees inferred under the three-, four-, five-, six- and seven-class group together crocodiles and birds with bootstrap supports of 90, 94, 99, 98 and 100 respectively (Figure 6A). The runtime of the one-class models (ModelFinder) and MixtureFinder on this dataset with 10 CPU threads were 0.24 hours and 1.35 hours respectively and memory requirement for those two analyses are 359 and 1295MB respectively.

**Figure 6.**
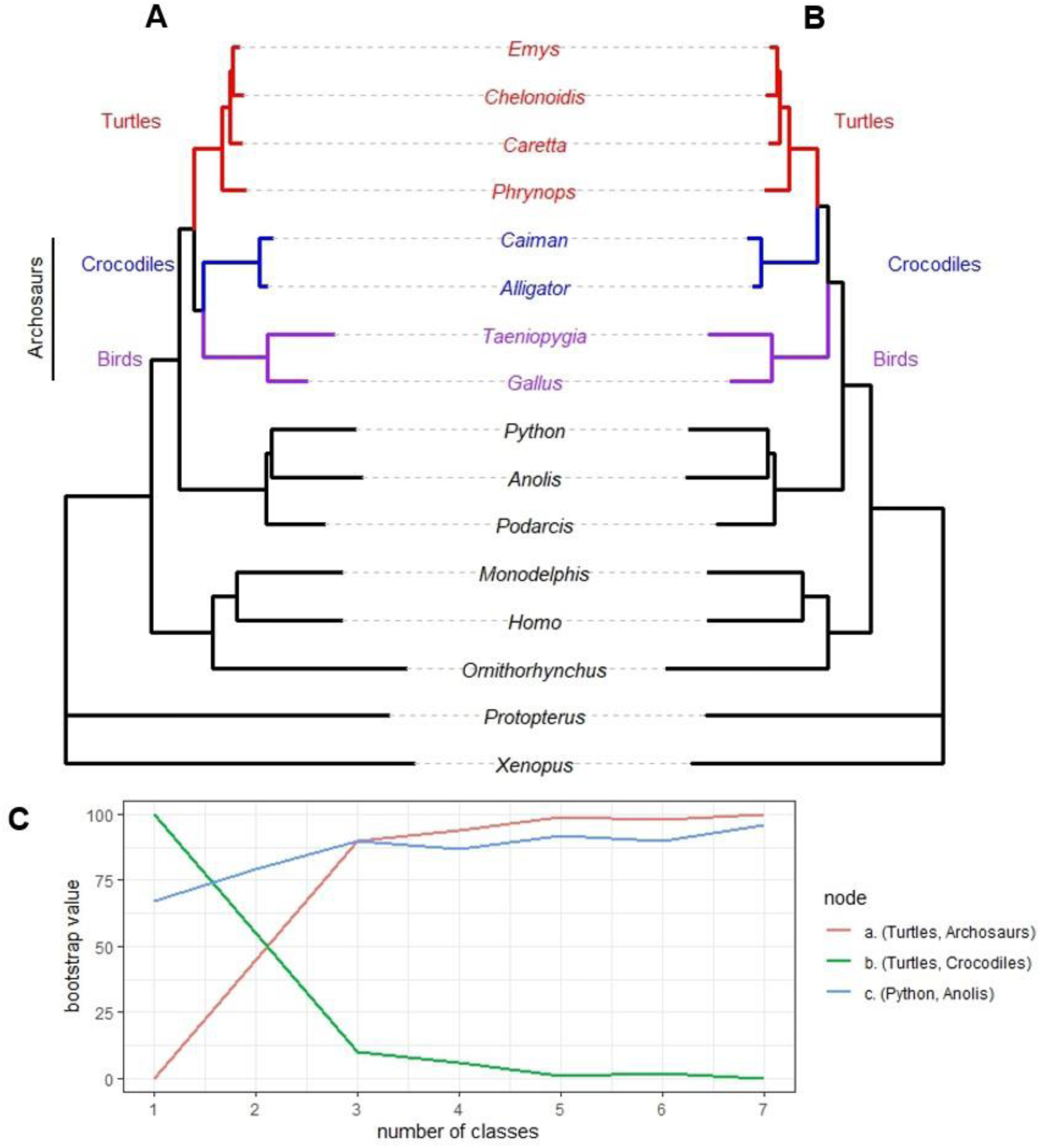
Mixture models change the conclusions from a concatenated phylogenomic dataset. **A.** The three-, four-, five-, six- and seven-class trees all recover a tree with Turtles as the sister group of Archosaurs with bootstrap supports of 90, 94, 99, 98 and 100 respectively. **B.** The one- and two-class trees both recover Turtles as the sister group of Crocodiles, with bootstrap supports of 100 and 55 respectively. **C.** Bootstrap supports of three key nodes in the tree (Turtles-Archosaurs), (Turtles-Crocodiles) and (Python-Anolis) change almost monotonically as classes are added to the mixture model. Bootstrap supports of the rest of the nodes in the trees are 100%.

## Discussion

In this study we introduce MixtureFinder, a simple algorithm implemented in IQ-TREE2 which can be used to estimate the optimal number of classes for mixture models of DNA sequence evolution. In so doing, we also test whether a commonly made assumption in phylogenetics: that the small data blocks used in partitioned models can be adequately modelled by a one-class substitution model. Surprisingly, we show that two-class mixture models of DNA sequence evolution almost always fit these small data blocks better than one class models, and that this appears to improve the estimated tree topologies and increase their lengths. We hypothesise that the increased tree lengths arise because the additional parameters in the two-class models allow them to infer more of the substitutions underlying the empirical alignments. This suggests that partitioned models may fail to adequately account for the complexity of molecular evolution in many phylogenomic studies.

In phylogenetic studies, researchers often use partitioned models to describe highly heterogeneous alignments. In these models it is typical to partition an alignment into the smallest possible units based on biological information about the sequences, such as defining each codon position from each protein coding gene as a single data block in a DNA sequence alignment (Shapiro et al. 2006), or defining each protein domain as a data block in an amino acid alignment (Zoller et al. 2015). Following this, some researchers assign an independent substitution model to each data block, while others merge similar data blocks into larger partitions using algorithms such as PartitionFinder and PartitionTest (Posada and Crandall 2001; Posada and Buckley 2004; Sullivan and Joyce 2005; Lanfear et al. 2012; Darriba and Posada 2015). Our study suggests that such approaches may limit the accuracy of phylogenetic inferences, because the single class model which is assigned to each partition is unlikely to adequately model the complex molecular evolutionary processes underlying that partition (Figure 2). In some ways this is surprising, because the single partitions we use in this study are usually quite small (e.g. just a few hundred base pairs), and so contain limited information. However, in other ways we should not be surprised. It has been appreciated for some time that even the most complex of the commonly used stationary and reversible substitution models in phylogenetics, the GTR model, represents a huge oversimplification of the complex molecular evolutionary processes that underly most alignments (Gatesy 2007). Our results showing that most individual partitions are better fit by two-rather than one-class models support the view that one class models tend to be too simple, even for short alignments.

This study extends the practical use of mixtures of Q matrices from amino acid alignments to DNA sequence alignments. Phylogenetic studies of amino acids routinely use complex mixture models. These include mixtures of Q matrices such as UL2, UL3, EXEHO, LG4M, LG4X, etc. (Le et al. 2008; Wang et al. 2008; Le et al. 2012), and mixtures of amino acid frequencies such as the CAT and PMSF models (Wang et al. 2018), which imply mixtures of Q matrices. In most cases the parameters of amino acid mixture models are estimated in advance on very large datasets, and they are then applied to empirical datasets with fixed parameters. This is done because the computational demands of estimating amino acid mixture models are high, and the complexity of the models mean that accurate parameter estimates typically require very large datasets beyond the size of many empirical datasets. In this study, we have shown that estimating mixtures of Q matrices for DNA sequence alignments is practical and the parameters of the underlying models (including the number of classes, and the model parameters of each class) can be quite close to the actual values on the simulated datasets, even when the size of the dataset is modest relative to those commonly used in empirical studies (e.g. Figure 4). Because of this, it will often be practical for empiricists to estimate DNA mixture models for their own alignments, and to then use these models for tree inference. However, it still important to note that the computational demands of estimating mixtures of Q matrices for DNA sequence alignments are much higher than the standard DNA substitution models, and the differences in topologies that we recorded were often marginal. Because of this, the benefits of applying Q mixture models for the specific empirical analyses need to be carefully weighed against the additional computational demands. Mixtures of Q matrices are most likely to offer meaningful gains for uncertain nodes in a phylogeny, for example those for which different studies have recovered different results (as in our example in Figure 6), or those where there is a great of underlying variation due to processes such as incomplete lineage sorting (e.g. the backbone of the phylogeny of birds Stiller et al. 2024). In the latter case, the slight improvements in gene trees expected from applying mixture models may prove useful in correctly resolving nodes in a phylogeny where evidence in support of different species trees is almost equivocal. These potential benefits should be weighed against the increased computational demands of applying such models.

Our application of MixtureFinder to a dataset of vertebrate taxa suggest that DNA mixture models may help to recover accurate tree topologies. The dataset we used was constructed to examine the phylogenetic relationships among turtles, crocodiles, birds, and other vertebrates (Chiari et al. 2012). This has been a relatively contentious area of the tree of life, with many studies, including the most recent, suggesting that turtles are the sister group to a clade comprising birds and crocodiles (the archosaurs: Iwabe et al. 2005; Chiari et al. 2012; Crawford et al. 2012; Fong et al. 2012; Lee 2013; Field et al. 2014), while other studies have reached different conclusions (Meyer and Zardoya 2003; Lyson et al. 2012). The dataset we used in this study is notable because it has already been shown that dramatically over-simplified models (e.g. unpartitioned models) tend to recover trees which disagree with the modern consensus (that turtles are the sister group of archosaurs), while more complex (e.g. highly partitioned) models recover strong support for trees that agree with it, and that such partitioned models tend be strongly preferred using standard model selection methods. Our results using MixtureFinder recapitulate these results and show a remarkable pattern. We show that a one-class model with the concatenated alignment (i.e. an unpartitioned, non-mixture model), recovers a clade comprising turtles and crocodiles with high bootstrap support (100%, Figure 6C). However, as we progressively add classes to the mixture model, the support for this clade decreases, while the support for the turtles as sister to the archosaurs increases (Figure 6C). Under the two-class mixture model the maximum likleihood tree still supports turtles and crocodiles as sister groups, but with low bootstrap support (55%). But under the three-, four-five-six- and seven-class models, the best tree supports turtles as the sister group to archosaurs, with increasing bootstrap support for the latter group (90%, 94%, 99%, 98% and 100% respectively). MixtureFinder suggests that the best model is the seven-class model. The conclusions using mixture models thus match the modern consensus that turtles are the sister group to archosaurs and give another example of the sensitivity of some nodes in the tree of life to the assumed model of sequence evolution.

Our analyses suggest model selection with mixture models is relatively insensitive to the choice of model selection criterion. Besides the frequently-relatively insensitive AIC and BIC, we also implemented the Likelihood Ratio Test (LRT) for model selection. One issue with this is in our implementation of the LRT we assume that the Likelihood Ratio Statistic (LRS) follows a chi-square distribution but this is known not to be the case for mixture models (Mitchell et al. 2019). The true distribution of the LRS remains unexplored for the case we uses here. To test whether our implemented LRT is reasonable, we showed the results on empirical data (Supplementary Figure S2), which gave similar results to BIC, and simulation data (Supplementary Figure S9), which is close to the LRS threshold under the chi-square distribution when the degrees of freedom for added Q matrix are small. However, because the potential weakness of the LRT that LRS for mixture hypotheses do not have chi-squared distributions (Self and Liang 1987; Mitchell et al. 2019), and because the BIC and AIC remain the most commonly used model selection criteria in phylogenetics, we use BIC as the default option in MixtureFinder. We acknowledge that these approaches may also have limitations (Susko and Roger 2020), although our simulations suggest that they perform acceptably in a wide range of cases.

One current frustration in phylogenetic model selection is that we know of no objective method to compare the fit of mixture models to partitioned models to a given alignment (Crotty and Holland 2022). This means that empiricists will still need to make a reasoned choice as to the best type of model to use for a particular dataset, and when mixture models and partitioned models give differing conclusions, it may be difficult to rule out either result. However, given the overwhelming evidence that a mixture model almost always fits individual partitions better than a one-class model, it seems that the conclusions suggested by using mixture models on DNA sequence alignments are certainly worth considering.

## Materials and Methods

### Empirical datasets for testing two-class models vs one-class models

We selected three large and well-annotated DNA sequence alignments of protein coding genes to compare the fit of one- and two-class models, shown in table two. We assigned each codon position in each gene to a separate partition, resulting in 15486 partitions from mammals, 424 from metazoans (first and second codon positions only, the original study removed third codon positions), and 3924 from plants.

**Table 2.** Empirical datasets used in the test.

|  | Species | Taxa | Genes | DNA<br>Partitions | Reference |
| --- | --- | --- | --- | --- | --- |
| **A** | Mammal | 90 | 5162 | 15486 | Wu et al. 2018 |
| **B** | Metazoan | 78 | 212 | 424 | Cannon et al. 2016 |
| **C** | Seed plant | 38 | 1308 | 3924 | Ran et al. 2018 |

For each partition, we used ModelFinder (Kalyaanamoorthy et al. 2017) to select the best one-class model, and MixtureFinder (up to two classes) to select the best two-class model. We then compared the BIC values or applied the LRT to determine the best model. Using these models, we then estimated the maximum likelihood tree for each partition using the one-class model (which we call the one-class tree), and the maximum likelihood tree using the two-class model (which we call the two-class tree).

We compared the one-class and two-class trees to estimates of the species tree obtained via concatenated ML analysis of amino acid alignments done in previous studies. The concatenated amnio acid tree for the mammal dataset tree was taken from Naser-Khdour et al. (2022) which was calculated using a partitioned model. The concatenated ML trees for the metazoan and plant datasets were taken from the original publications listed in table 1 (both of which used an LG+I+G model on the concatenated amino acid alignments). To compare the one- and two-class partitioned trees to these datasets, we calculated the normalised Robinson-Foulds distance, which is simply the standard Robison-Foulds distance (Robinson and Foulds 1981) divided by the maximum possible Robinson-Foulds distance for the pair of trees being compared. Thus, the nRF is zero when the two trees being compared have identical topologies, and 1 if the two trees being compared have no splits in common.

### Testing MixtureFinder by simulation

We tested the ability of the MixtureFinder algorithm to recover mixture models with known numbers of classes. Using AliSim (Ly-Trong et al. 2022), we simulated mixture models with 1 to 5 classes on alignments of 1000 bp, 2000 bp, 5000 bp and 10000 bp with 100 taxa. Each class in a mixture model has an associated weight. We simulated class weights for each simulation under a uniform distribution between 0.1 and 1 and then normalised them so that they sum to 1. We repeated each combination of simulation conditions 20 times, for a total of 400 simulations. We used two different schemes to simulate the parameters of transition rates, base frequencies, RHAS models and tree lengths. First (called scheme 1 in this research), we followed a scheme similar to that of Wong et al. (2024) which simulates models using random numbers generated from plausible ranges. This tends to create simulated mixture models in which the models for each class are very different from one another. In this scheme, transition rates were simulated as six random integers selected between 5 and 50, then divided by the value of G↔T; base frequencies were simulated from a uniform distribution between 1 and 10 and normalised to sum to 1; and RHAS was simulated by Gamma model with alpha parameter under an exponential distribution that the mean equalled 1. The tree topology was simulated as Yule-Harding tree shape by AliSim and the branch length fitted an exponential distribution that the mean equalled 0.1.

Second (called scheme 2 in this research), we used parameters estimated from large collections of biological datasets (https://github.com/Cibiv/EvoNAPS), which fit one-class substitution models with invariant sites and gamma rate parameters (+I+G) to a large range of empirical datasets. This approach tends to simulate mixture models in which the models for each class are much more similar to each other, representing a much more challenging task for an algorithm to determine the underlying number of classes in the mixture. These models certainly represent real empirical datasets better than the first simulation scheme above, though we note that they may underestimate the true variability between models if the underlying models are actually mixture models. The EvoNAPS database contains tens of thousands of models of sequence evolution, from the most complex GTR model down to the simplest JC model. To select parameters from this set of models we first used the method of Naser-Khdour et al. (2021) to estimate the distribution of each of the transition rates and base frequencies, for each type of model (e.gg. GTR, HKY, JC). We then used the complete set of models and the method of Naser-Khdour et al. (2021) to estimate the distributions of the I+G model parameters and the lengths of the internal and external branches. All these distributions are shown in Supplementary Table S1 and S2. We then used a two-step process to select a model with which to simulate alignments. First, we selected the type of model (e.g. GTR, HKY, JC) using the distribution model types in the EvoNAPS database. Then, we generated random parameters from the appropriate parameter distributions above (e.g. GTR parameters are generated from the GTR parameter distributions), and the +I+G and branch length distributions from the global distributions generated above. The tree topology was simulated as Yule-Harding tree shape using AliSim. As above, we simulated datasets with 100 taxa, at lengths of 1000 bp, 2000 bp, 5000 bp and 10,000 bp, and we performed 20 replicates of each set of simulation conditions.

We additionally ran Pythia (Haag et al. 2022) on the simulated alignments and the distributions of the Pythia-based difficulty for phylogenetic tree inference in Supplementary Figure S10. Most of our simulated alignments had difficulty scores greater than 0.2, whereas more than 40% of the empirical alignments used to train Pythia, obtained from TreeBASE (Piel et al. 2000), had scores below 0.1. This indicates that our simulated alignments are more “challenging” than the empirical alignments. Additionally, alignments simulated under scheme 2 had slightly higher difficulty scores than those from scheme 1. The difficulty scores were also correlated with alignment length, with the shortest alignments (1000 bp) showing significantly higher difficulty than longer alignments in both simulation schemes.

Measuring the accuracy of estimated model parameters can be challenging for mixture models, because the estimated model may have a different number of classes to the simulated model. To address this, we use the recently-proposed Integrated Squared Error (ISE, Baños et al. 2024). This approach compares the cumulative density functions (CDF) of model parameters based on the weights of the mixture classes. In our application we first calculated the CDF of the transition rates and base frequencies for both the simulated and estimated models for each alignment, and then use these to calculate the integrated squared error (ISE) of the estimated model following.

### Testing the accuracy of LRT as model selection criteria by simulation

The chi-squared test for LRT in the phylogenetic model selection is not widely used. Particularly for mixture models, the chi-squared distribution does not fit the hypothesis (Mitchell et al. 2019). To address one potential limitation, we tested the accuracy the LRT that we implemented using simulations under the null model. To do this, we simulated 1,000 datasets under a 1-class model based on simulation scheme 2 that described above, including 250 alignments each with 1000 bp, 2000 bp, 5000 bp and 10000 bp. We ran MixtureFinder on these alignments up to a 2-class model and calculated the Likelihood Ratio Statistics, LRS = 2*(lnL_2_ – lnL_1_), comparing the 2-class and 1-class model. We then compared the results of using the 95^th^ percentile of LRS and LRT with a p-value threshold 0.05.

### Analysis of the empirical dataset of vertebrates

We used IQ-TREE2 to analyse a vertebrate dataset containing 16 taxa and 248 genes, with a total length of 186 Kbp (Chiari et al. 2012). First, we ran MixtureFinder using IQ-TREE2 version 2.3.5.2.mf to estimate the best mixture model (using only GTR models for each mixture class), including estimating the number of classes in the model, and estimating the ML tree using this model with 1000 ultrafast bootstraps. This analysis suggested that the best number of mixture components was seven. We then ran six additional analyses estimating the ML tree with 1000 ultrafast bootstraps using 1, 2, 3, 4, 5 and 6 GTR model classes. All command lines for these analyses are provided in the Supplementary Text 4.

## Data and Resource Availability

The corresponding IQ-TREE2 software version is freely available at http://www.iqtree.org/. The three partitioned empirical datasets can be download at https://github.com/roblanf/BenchmarkAlignments. The raw data for the figures, simulated MSAs (with python scripts written to generate it), and the vertebrate alignments (with the IQ-TREE2 files that infer the one- to seven-class trees) are available at https://figshare.com/articles/dataset/MixtureFinder_logs/24463921.

## Acknowledgements

This work was supported by an Australian Research Council Discovery Project (DP200103151); a Moore-Simons Foundation grant (https://doi.org/10.46714/735923LPI to B.Q.M.); and a Chan-Zuckerberg Initiative Grant for Essential Open-Source Software for Science to B.Q.M. and R.L. We thank Nhan Ly-Trong, Caitlin Cherryh, Frederick Jaya, Jeremias Ivan and Matthew Macaulay for their comments and discussion of this work.

## Supplementary Materials

The Supplementary Text, Tables and Figures are available at [supplementary file link].

## Notes

### Competing Interest Statement

The authors have declared no competing interest.

### Summary of Updates

Updating the results and discussion of the Likelihood Ratio Test

## Reference

Abadi, S., Azouri, D., Pupko, T. and Mayrose, I. 2019. Model selection may not be a mandatory step for phylogeny reconstruction. Nature communications 10(1), p. 934.

Akaike, H. 1974. A new look at the statistical model identification. In: Springer Series in Statistics. Springer series in statistics. New York, NY: Springer New York, pp. 215–222.

Anderson, F.E. and Swofford, D.L. 2004. Should we be worried about long-branch attraction in real data sets? Investigations using metazoan 18S rDNA. Molecular phylogenetics and evolution 33(2), pp. 440–451.

Baños, H., Susko, E. and Roger, A.J. 2024. Is over-parameterization a problem for profile mixture models? Systematic biology 73(1), pp. 53–75.

Bejerano, G., Pheasant, M., Makunin, I., Stephen, S., Kent, W.J., Mattick, J.S. and Haussler, D. 2004. Ultraconserved elements in the human genome. Science (New York, N.Y.) 304(5675), pp. 1321–1325.

Blanquart, S. and Lartillot, N. 2008. A site- and time-heterogeneous model of amino acid replacement. Molecular biology and evolution 25(5), pp. 842–858.

Brandley, M.C., Schmitz, A. and Reeder, T.W. 2005. Partitioned Bayesian analyses, partition choice, and the phylogenetic relationships of scincid lizards. Systematic biology 54(3), pp. 373– 390.

Cannon, J.T., Vellutini, B.C., Smith, J., Ronquist, F., Jondelius, U. and Hejnol, A. 2016. Xenacoelomorpha is the sister group to Nephrozoa. Nature 530(7588), pp. 89–93.

Chiari, Y., Cahais, V., Galtier, N. and Delsuc, F. 2012. Phylogenomic analyses support the position of turtles as the sister group of birds and crocodiles (Archosauria). BMC biology 10(1), p. 65.

Crawford, N.G., Faircloth, B.C., McCormack, J.E., Brumfield, R.T., Winker, K. and Glenn, T.C. 2012. More than 1000 ultraconserved elements provide evidence that turtles are the sister group of archosaurs. Biology letters 8(5), pp. 783–786.

Crotty, S.M. and Holland, B.R. 2022. Comparing partitioned models to mixture models: Do information criteria apply? Systematic biology 71(6), pp. 1541–1548.

Crotty, S.M., Minh, B.Q., Bean, N.G., Holland, B.R., Tuke, J., Jermiin, L.S. and Haeseler, A.V. 2020. GHOST: Recovering historical signal from heterotachously evolved sequence alignments. Systematic biology 69(2), pp. 249–264.

Crotty, S.M., Rohrlach, A.B., Ndunguru, J. and Boykin, L.M. 2018. Characterising genetic diversity in cassava Brown Streak Virus. bioRxiv. Available at: 10.1101/455303.

Darriba, D. and Posada, D. 2015. The impact of partitioning on phylogenomic accuracy. bioRxiv. Available at: 10.1101/023978.

Felsenstein, J. 2002. Inferring Phylogenies. New York, NY: Oxford University Press.

Field, D.J., Gauthier, J.A., King, B.L., Pisani, D., Lyson, T.R. and Peterson, K.J. 2014. Toward consilience in reptile phylogeny: miRNAs support an archosaur, not lepidosaur, affinity for turtles. Evolution & development 16(4), pp. 189–196.

Fletcher, R. 2013. Practical methods of optimization. 2nd ed. Chichester, England: John Wiley & Sons.

Fong, J.J., Brown, J.M., Fujita, M.K. and Boussau, B. 2012. A phylogenomic approach to vertebrate phylogeny supports a turtle-archosaur affinity and a possible paraphyletic lissamphibia. PloS one 7(11), p. e48990.

Garcia-España, A., Mares, R., Sun, T.-T. and Desalle, R. 2009. Intron evolution: testing hypotheses of intron evolution using the phylogenomics of tetraspanins. PloS one 4(3), p. e4680.

Gatesy, J. 2007. A tenth crucial question regarding model use in phylogenetics. Trends in ecology & evolution 22(10), pp. 509–510.

Gelman, A., Carlin, J.B., Stern, H.S. and Rubin, D.B. 1995. Bayesian Data Analysis. Philadelphia, PA: Chapman & Hall/CRC.

Guimarães Fabreti, L. and Höhna, S. 2023. Nucleotide substitution model selection is not necessary for Bayesian inference of phylogeny with well-behaved priors. Systematic biology 72(6), pp. 1418–1432.

Haag, J., Höhler, D., Bettisworth, B. and Stamatakis, A. 2022. From easy to hopeless-predicting the difficulty of phylogenetic analyses. Molecular biology and evolution 39(12). Available at: 10.1093/molbev/msac254.

Hasegawa, M., Kishino, H. and Yano, T. 1985. Dating of the human-ape splitting by a molecular clock of mitochondrial DNA. Journal of molecular evolution 22(2), pp. 160–174.

Iwabe, N., Hara, Y., Kumazawa, Y., Shibamoto, K., Saito, Y., Miyata, T. and Katoh, K. 2005. Sister group relationship of turtles to the bird-crocodilian clade revealed by nuclear DNA-coded proteins. Molecular biology and evolution 22(4), pp. 810–813.

Jukes, T.H. and Cantor, C.R. 1969. Evolution of Protein Molecules. In: Mammalian Protein Metabolism. Elsevier, pp. 21–132.

Kainer, D. and Lanfear, R. 2015. The effects of partitioning on phylogenetic inference. Molecular biology and evolution 32(6), pp. 1611–1627.

Kalyaanamoorthy, S., Minh, B.Q., Wong, T.K.F., von Haeseler, A. and Jermiin, L.S. 2017. ModelFinder: fast model selection for accurate phylogenetic estimates. Nature methods 14(6), pp. 587–589.

Kapli, P., Kotari, I., Telford, M.J., Goldman, N. and Yang, Z. 2023. DNA sequences are as useful as protein sequences for inferring deep phylogenies. Systematic biology 72(5), pp. 1119–1135.

Kolaczkowski, B. and Thornton, J.W. 2008. A mixed branch length model of heterotachy improves phylogenetic accuracy. Molecular biology and evolution 25(6), pp. 1054–1066.

Lanfear, R., Calcott, B., Ho, S.Y.W. and Guindon, S. 2012. Partitionfinder: combined selection of partitioning schemes and substitution models for phylogenetic analyses. Molecular biology and evolution 29(6), pp. 1695–1701.

Lanfear, R., Frandsen, P.B., Wright, A.M., Senfeld, T. and Calcott, B. 2016. PartitionFinder 2: New methods for selecting partitioned models of evolution for molecular and morphological phylogenetic analyses. Molecular biology and evolution, p. msw260.

Lartillot, N. and Philippe, H. 2004. A Bayesian mixture model for across-site heterogeneities in the amino-acid replacement process. Molecular biology and evolution 21(6), pp. 1095–1109.

Le, S.Q., Dang, C.C. and Gascuel, O. 2012. Modeling protein evolution with several amino acid replacement matrices depending on site rates. Molecular biology and evolution 29(10), pp. 2921–2936.

Le, S.Q., Lartillot, N. and Gascuel, O. 2008. Phylogenetic mixture models for proteins. Philosophical transactions of the Royal Society of London. Series B, Biological sciences 363(1512), pp. 3965–3976.

Lee, M.S.Y. 2013. Turtle origins: insights from phylogenetic retrofitting and molecular scaffolds. Journal of evolutionary biology 26(12), pp. 2729–2738.

Lemmon, A.R., Emme, S.A. and Lemmon, E.M. 2012. Anchored hybrid enrichment for massively high-throughput phylogenomics. Systematic biology 61(5), pp. 727–744.

Lyson, T.R., Sperling, E.A., Heimberg, A.M., Gauthier, J.A., King, B.L. and Peterson, K.J. 2012. MicroRNAs support a turtle + lizard clade. Biology letters 8(1), pp. 104–107.

Ly-Trong, N., Naser-Khdour, S., Lanfear, R. and Minh, B.Q. 2022. AliSim: A fast and versatile phylogenetic sequence simulator for the genomic era. Molecular biology and evolution 39(5). Available at: 10.1093/molbev/msac092.

Meyer, A. and Zardoya, R. 2003. Recent advances in the (molecular) phylogeny of vertebrates. Annual review of ecology, evolution, and systematics 34(1), pp. 311–338.

Minh, B.Q., Schmidt, H.A., Chernomor, O., Schrempf, D., Woodhams, M.D., von Haeseler, A. and Lanfear, R. 2020. IQ-TREE 2: New models and efficient methods for phylogenetic inference in the genomic era. Molecular biology and evolution 37(5), pp. 1530–1534.

Mitchell, J.D., Allman, E.S. and Rhodes, J.A. 2019. Hypothesis testing near singularities and boundaries. Electronic journal of statistics 13(1), pp. 2150–2193.

Naser-Khdour, S., Minh, B.Q. and Lanfear, R. 2021. The influence of model violation on phylogenetic inference: A simulation study. bioRxiv. Available at: 10.1101/2021.09.22.461455.

Naser-Khdour, S., Quang Minh, B. and Lanfear, R. 2022. Assessing confidence in root placement on phylogenies: an empirical study using nonreversible models for mammals. Systematic Biology 71(4), pp. 959–972.

Nylander, J.A.A., Ronquist, F., Huelsenbeck, J.P. and Nieves-Aldrey, J.L. 2004. Bayesian phylogenetic analysis of combined data. Systematic biology 53(1), pp. 47–67.

Pagel, M. and Meade, A. 2004. A phylogenetic mixture model for detecting pattern-heterogeneity in gene sequence or character-state data. Systematic biology 53(4), pp. 571–581.

Piel, W.H., Donoghue, M., Sanderson, M. and Netherlands, L. 2000.TreeBASE: a database of phylogenetic information.

Posada, D. and Buckley, T.R. 2004. Model selection and model averaging in phylogenetics: advantages of akaike information criterion and bayesian approaches over likelihood ratio tests. Systematic biology 53(5), pp. 793–808.

Posada, D. and Crandall, K.A. 2001. Selecting the best-fit model of nucleotide substitution. Systematic biology 50(4), pp. 580–601.

Quang, L.S., Gascuel, O. and Lartillot, N. 2008. Empirical profile mixture models for phylogenetic reconstruction. *Bioinformatics (Oxford*, England) 24(20), pp. 2317–2323.

Ran, J.-H., Shen, T.-T., Wang, M.-M. and Wang, X.-Q. 2018. Phylogenomics resolves the deep phylogeny of seed plants and indicates partial convergent or homoplastic evolution between Gnetales and angiosperms. *Proceedings*. Biological sciences 285(1881), p. 20181012.

Redmond, A.K. and McLysaght, A. 2021. Evidence for sponges as sister to all other animals from partitioned phylogenomics with mixture models and recoding. Nature communications 12(1), p. 1783.

Robinson, D.F. and Foulds, L.R. 1981. Comparison of phylogenetic trees. Mathematical biosciences 53(1–2), pp. 131–147.

Schrempf, D., Lartillot, N. and Szöllősi, G. 2020. Scalable empirical mixture models that account for across-site compositional heterogeneity. Molecular biology and evolution 37(12), pp. 3616–3631.

Schwarz, G. 1978. Estimating the dimension of a model. Annals of statistics 6(2), pp. 461–464.

Self, S.G. and Liang, K.-Y. 1987. Asymptotic properties of maximum likelihood estimators and likelihood ratio tests under nonstandard conditions. Journal of the American Statistical Association 82(398), pp. 605–610.

Shapiro, B., Rambaut, A. and Drummond, A.J. 2006. Choosing appropriate substitution models for the phylogenetic analysis of protein-coding sequences. Molecular biology and evolution 23(1), pp. 7–9.

Soubrier, J., Steel, M., Lee, M.S.Y., Der Sarkissian, C., Guindon, S., Ho, S.Y.W. and Cooper, A. 2012. The influence of rate heterogeneity among sites on the time dependence of molecular rates. Molecular biology and evolution 29(11), pp. 3345–3358.

Stiller, J. et al. 2024. Complexity of avian evolution revealed by family-level genomes. Nature 629(8013), pp. 851–860.

Sullivan, J. and Joyce, P. 2005. Model selection in phylogenetics. Annual review of ecology, evolution, and systematics 36(1), pp. 445–466.

Susko, E. and Roger, A.J. 2020. On the use of information criteria for model selection in phylogenetics. Molecular biology and evolution 37(2), pp. 549–562.

Tavaré, S. 1986. Some probabilistic and statistical problems on the analysis of DNA sequence. Lecture of Mathematics for Life Science 17, p. 57.

Wang, H.-C., Li, K., Susko, E. and Roger, A.J. 2008. A class frequency mixture model that adjusts for site-specific amino acid frequencies and improves inference of protein phylogeny. BMC evolutionary biology 8(1), p. 331.

Wang, H.-C., Minh, B.Q., Susko, E. and Roger, A.J. 2018. Modeling site heterogeneity with posterior mean site frequency profiles accelerates accurate phylogenomic estimation. Systematic biology 67(2), pp. 216–235.

Wang, H.-C., Susko, E. and Roger, A.J. 2019. The relative importance of modeling site pattern heterogeneity versus partition-wise heterotachy in phylogenomic inference. Systematic biology 68(6), pp. 1003–1019.

Whelan, N.V. and Halanych, K.M. 2016. Who let the CAT out of the bag? Accurately dealing with substitutional heterogeneity in phylogenomic analyses. *Systematic biology*, p. syw084.

Wong, T.K.F., Cherryh, C., Rodrigo, A.G., Hahn, M.W., Minh, B.Q. and Lanfear, R. 2024. MAST: Phylogenetic inference with mixtures across sites and trees. Systematic biology. Available at: 10.1093/sysbio/syae008.

Wu, S., Edwards, S. and Liu, L. 2018. Genome-scale DNA sequence data and the evolutionary history of placental mammals. Data in brief 18, pp. 1972–1975.

Yang, Z. 1994. Maximum likelihood phylogenetic estimation from DNA sequences with variable rates over sites: approximate methods. Journal of molecular evolution 39(3), pp. 306– 314.

Yang, Z. 1995. A space-time process model for the evolution of DNA sequences. Genetics 139(2), pp. 993–1005.

Yang, Z. 1996. Maximum-likelihood models for combined analyses of multiple sequence data. Journal of molecular evolution 42(5), pp. 587–596.

Zoller, S., Boskova, V. and Anisimova, M. 2015. Maximum-likelihood tree estimation using Codon substitution models with multiple partitions. Molecular biology and evolution 32(8), pp. 2208–2216.

