## Supplementary for "MixtureFinder: Estimating DNA mixture models for phylogenetic analyses"

**Supplementary Texts**

**1. IQ-TREE 2 command lines to test two-class models vs one-class models**

One-class model:

iqtree2 -s alignment.nex -m MF -mfreq FO

Two-class model:

iqtree2 -s alignment.nex -m MIX"{*Q1*,*Q2*}+*RHAS*" -te *1classtree.treefile*

where *Q1* is the Q matrix selected in the one-class model, *RHAS* is the rate heterogeneity among site model selected in the one-class model, *1classtree.treefile* the tree estimated by the one class model and *Q2* is the second Q matrix in the two-class model. (E.g., -m MIX"{GTR+FO,HKY+FO}+I+G")

Tree inference of the two-class model:

iqtree2 -s alignment.nex -m MIX"{*Q1{p1}*,*Q2{p2}*}+*RHAS{pr}*"

where *Q1* is the Q matrix selected in the one-class model, *Q2* is the best Q matrix selected in the two-class model and *RHAS* is the rate heterogeneity among site model selected in the one-class model. *p1*, *p2* and *pr* are the parameters of the *Q1*, *Q2* and *RHAS* that estimated in the two-class model. (E.g., -m MIX"{GTR{1/1.5/1.5/1/0.5}+FO{0.2/0.3/0.3/0.2},HKY{0.5}+FO{0.3/0.3/0.2/0.2}}+I{0.1}+G{0.5}")

**2. IQ-TREE 2 command lines to test class weight estimates when given true number of classes**

If the true number of classes is 1:

iqtree2 -s alignment.nex -m GTR+FO+I+G

If the true number of classes is greater than 1:

iqtree2 -s alignment.nex -m MIX"{*GTR+FO,…,GTR+FO*}+I+G"

where we input the true number of "*GTR+FO*" in the *MIX"{GTR+FO,…,GTR+FO}*".

**3. IQ-TREE 2 command lines to run MixtureFinder on simulated data**

We run the experiment on IQ-TREE 2 version 2.3.5.2.mf. Please note that the commands used for the mixture model differ from those that you should use in the latest version of IQ-TREE, because we have since updated the user interface. The appropriate commands for the latest version are given in the tutorial http://www.iqtree.org/doc/Complex-Models#mixture-models.

The following analyses run the MixtureFinder algorithm, to estimate the best number of mixture classes, the best model of the distribution of rates across sites (after the best number of classes is estimated), and the maximum likelihood tree under the best mode. The analysis uses BIC to estimate the number of classes.

Q-mixture:

```
iqtree2 -s alignment.nex -m MIX+MFP -mrate E,I,G,I+G,R,I+R
```

and GTR-mixture:

```
iqtree2 -s alignment.nex -m MIX+MFP -mset GTR -mrate E,I,G,I+G,R,I+R
```

**4. IQ-TREE 2 command lines for analysing the vertebrate dataset**

4.1 MixtureFinder analysis

The following analysis runs the MixtureFinder algorithm, to estimate the best number of mixture classes (assuming each class is a GTR model), and the maximum likelihood tree under the best model with 1000 ultrafast bootstraps. The analysis uses the BIC for all aspects of model selection, including selecting the best number of mixture classes. We run the experiment on IQ-TREE 2 version 2.3.5.2.mf.

iqtree2 -s alignment.nex -m MIX+MFP -mset GTR -mrate E,I,G,I+G,R,I+R -bb 1000

4.2 One- to six-class models and trees

The following command lines estimate a one class model (the first line) and then mixture models with 2-6 GTR model classes. Each analysis uses the Q matrices parameters and rate heterogenous model from the IQ-TREE 2 model checkpoint file from section 4.1, and all include 1000 ultrafast bootstraps.

iqtree2 –s alignment.nex -m GTR+FO+I+R3 -bb 1000
iqtree2 –s alignment.nex -m MIX"{GTR+FO,…,GTR+FO}+I+R3" -bb 1000

where we input the true number of "*GTR+FO*" in the *MIX"{GTR+FO,…,GTR+FO}"*.

**Supplementary Tables**

**Table S1 Distribution of parameters in substitution models in simulation scheme 2.**

| Substitution model | Frequencies* | Parameter | Distribution | Shape | Location | Scale |
|---|---|---|---|---|---|---|
| F81 | 0.0009 | $F_A$ | Log Gamma | 0.072 | 0.345 | 0.005 |
| | | $F_C$ | Generalized Exponential | 0.420, 3.653, 1.379 | 0.140 | 0.189 |
| | | $F_G$ | Generalized Logistic | 0.437 | 0.285 | 0.017 |
| | | $F_T$ | Exponential Normal | 2982 | 0.154 | 2.234e-05 |
| GTR | 0.1133 | A↔C | Inverse Weibull | 3.386 | -1.124 | 2.543 |
| | | A↔G | Alpha | 4.672 | -3.414 | 37.48 |
| | | A↔T | Exponential Weibull | 262.7, 0.357 | -0.177 | 0.008 |
| | | C↔G | Inverse Weibull | 3.580 | -1.244 | 2.431 |
| | | C↔T | Alpha | 3.914 | -4.390 | 38.77 |
| | | $F_A$ | Exponential Weibull | 0.583, 4.730 | 0.077 | 0.246 |
| | | $F_C$ | Maxwell | / | 0.083 | 0.098 |
| | | $F_G$ | Log Normal | 0.007 | -7.039 | 7.300 |
| | | $F_T$ | Exponential Normal | 0.893 | 0.191 | 0.040 |
| HKY | 0.400 | A↔G | Normal | / | 4.275 | 1.861 |
| | | C↔T | A↔G | | | |
| | | $F_A$ | Maxwell | / | 0.106 | 0.084 |
| | | $F_C$ | Exponential Power | 3.057 | 0.083 | 0.250 |
| | | $F_G$ | Log Normal | 0.007 | -7.039 | 7.300 |
| | | $F_T$ | Exponential Normal | 0.893 | 0.191 | 0.040 |
| JC | 0.0004 | / | | | | |
| K2P | 0.0567 | A↔G | Generalized Gamma | 0.653, 2.560 | 1.203 | 5.056 |
| | | C↔T | A↔G | | | |
| K3P | 0.0343 | A↔G | Double Weibull | 0.990 | 4.707 | 1.078 |
| | | A↔T | Alpha | 3.775 | -0.586 | 5.468 |
| | | C↔G | A↔T | | | |
| | | C↔T | A↔G | | | |
| K3Pu | 0.0254 | A↔G | Laplace | / | 4.089 | 1.000 |
| | | A↔T | Log Laplace | 3.778 | -0.025 | 0.781 |
| | | C↔G | A↔T | | | |
| | | C↔T | A↔G | | | |
| | | $F_A$ | Generalized Logistic | 3.288 | 0.163 | 0.043 |
| | | $F_C$ | Power Normal | 5.785 | 0.400 | 0.086 |
| | | $F_G$ | Generalized Logistic | 0.554 | 0.295 | 0.020 |
| | | $F_T$ | Exponential Normal | 1.746 | 0.163 | 0.023 |
| SYM | 0.0614 | A↔C | Log Laplace | 2.621 | 0.377 | 1.393 |
| | | A↔G | Log Laplace | 3.929 | -0.023 | 5.833 |
| | | A↔T | Inverse Weibull | 3.535 | -0.630 | 1.400 |

| | | | | | | |
|---|---|---|---|---|---|---|
| | | C↔G | Alpha | 3.261 | -0.617 | 6.378 |
| | | C↔T | Generalized Logistic | 651.7 | -10.14 | 2.455 |
| TIM2e | 0.0245 | A↔C | Log Gamma | 1368 | -141.2 | 19.76 |
| | | A↔G | Chi | 1.779 | 1.277 | 3.921 |
| | | A↔T | A↔C | | | |
| | | C↔T | Beta | 1.987, 10.20 | 1.200 | 37.43 |
| TIM3e | 0.0756 | A↔C | Exponential Normal | 2.326 | 1.273 | 0.258 |
| | | A↔G | Log Laplace | 4.130 | -1.546 | 7.771 |
| | | C↔G | A↔C | | | |
| | | C↔T | Log Normal | 0.522 | -0.189 | 7.15 |
| TIMe | 0.0175 | A↔G | Double Weibull | 1.099 | 4.524 | 0.867 |
| | | A↔T | Alpha | 2.780 | -0.192 | 2.876 |
| | | C↔G | A↔T | | | |
| | | C↔T | Double Gamma | 1.107 | 5.300 | 1.327 |
| TNe | 0.0275 | A↔G | Generalized Normal | 1.538 | 4.791 | 1.998 |
| | | C↔T | Generalized Gamma | 0.607, 2.895 | 1.022 | 7.242 |
| TPM2 | 0.0497 | A↔C | Double Gamma | 0.888 | 1.486 | 0.508 |
| | | A↔G | Beta | 1.740, 1.108e13 | 1.515 | 2.780e13 |
| | | A↔T | A↔C | | | |
| | | C↔T | A↔G | | | |
| TPM2u+F | 0.0489 | A↔C | Double Weibull | 0.953 | 1.478 | 0.350 |
| | | A↔G | Exponential Normal | 1.689 | 3.421 | 1.197 |
| | | A↔T | A↔C | | | |
| | | C↔T | A↔G | | | |
| | | $F_A$ | Inverse Weibull | 9200 | -416.6 | -416.8 |
| | | $F_C$ | Generalized Logistic | 0.298 | 0.333 | 0.014 |
| | | $F_G$ | Power Log Normal | 23.34, 0.038 | -2.013 | 2.468 |
| | | $F_T$ | Exponential Normal | 1.764 | 0.172 | 0.021 |
| TPM3 | 0.1317 | A↔C | Exponential Normal | 2.298 | 1.230 | 0.231 |
| | | A↔G | Minimum Extreme Value Weibull | 1.733 | 1.462 | 4.964 |
| | | C↔G | A↔C | | | |
| | | C↔T | A↔G | | | |
| TPM3u+F | 0.1057 | A↔C | Log Laplace | 4.374 | -0.399 | 2.232 |
| | | A↔G | Exponential Normal | 7.455 | 2.198 | 0.302 |
| | | C↔G | A↔C | | | |
| | | C↔T | A↔G | | | |
| | | $F_A$ | Exponential Power | 2.332 | 0.093 | 0.231 |
| | | $F_C$ | Inverse Weibull | 4.934e8 | -2.514e7 | 2.514e7 |
| | | $F_G$ | Inverse Gaussian | 0.018 | -0.052 | 17.02 |
| | | $F_T$ | Double Weibull | 1.323 | 0.247 | 0.042 |
| TVM+F | 0.0839 | A↔C | Inverse Weibull | 2.850 | -0.597 | 2.073 |
| | | A↔G | Alpha | 4.532 | -2.428 | 31.32 |

| | | | | | | |
|---|---|---|---|---|---|---|
| | | A↔T | Generalized Gamma | 12.90, 0.387 | 0.093 | 0.002 |
| | | C↔G | Inverse Gamma | 3.122 | -0.179 | 4.804 |
| | | C↔T | A↔G | | | |
| | | $F_A$ | Generalized Normal | 3.264 | 0.258 | 0.105 |
| | | $F_C$ | Double Gamma | 1.850 | 0.248 | 0.029 |
| | | $F_G$ | Maximum Extreme Value Weibull | 4.274 | 0.439 | 0.200 |
| | | $F_T$ | Power Normal | 0.179 | 0.176 | 0.028 |
| TVMe | 0.1025 | A↔C | Alpha | 3.776 | -0.914 | 10.30 |
| | | A↔G | Alpha | 5.032 | -4.409 | 50.12 |
| | | A↔T | Inverse Weibull | 3.146 | -0.521 | 1.362 |
| | | C↔G | Alpha | 2.912 | -0.582 | 5.774 |
| | | C↔T | A↔G | | | |

(* The frequency of each substitution model appears in the simulation, this also comes from the evoNAPS empirical data)

**Table S2 Distribution of RHAS model and tree branch length in simulation scheme 2.**

| Parameter | Distribution | Shape | Location | Scale |
|---|---|---|---|---|
| Invariable sites % | Generalized Pareto | -0.425 | -1.346e-10 | 0.345 |
| Gamma shape | Inverse Weibull | 5.576 | -1.346 | 2.408 |
| External branch | Alpha | 3.091e-11 | -0.006 | 0.012 |
| Internal branch | Alpha | 1.136e-09 | -0.003 | 0.007 |

**Supplementary Figures**

**Figure S1 MixtureFinder model initialisation strategy for k-class models (k >1).** Each candidate mixture model is initialised from the nested model with the highest likelihood. To ensure that the likelihood of the k-class model is not lower than that of the (k-1)-class model, the initial weight of the k[th] class ($W_k$) will iteratively reduce when the likelihood of k-class model is lower, up to 5 times. Meanwhile, to ensure that optimisation of each model works well, the model with low initial $W_k$ cannot be restored for further initialisation.

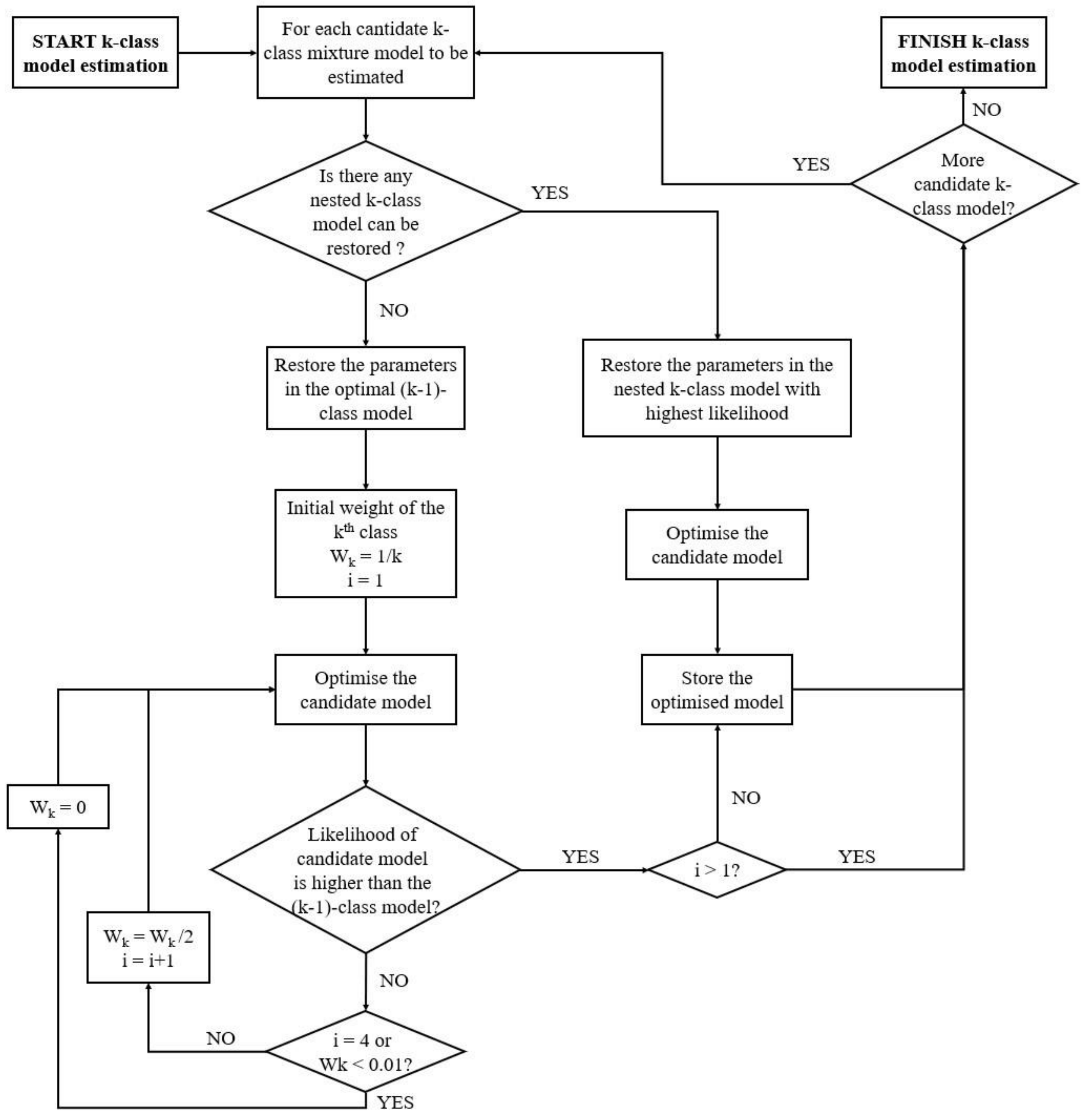

**Figure S2 The proportion of partitions in which the 1- or 2-class model are better according to likelihood ratio test (LRT).** The exact number of partitions in which the 1- or 2-class model are better is shown on the top of each bar.

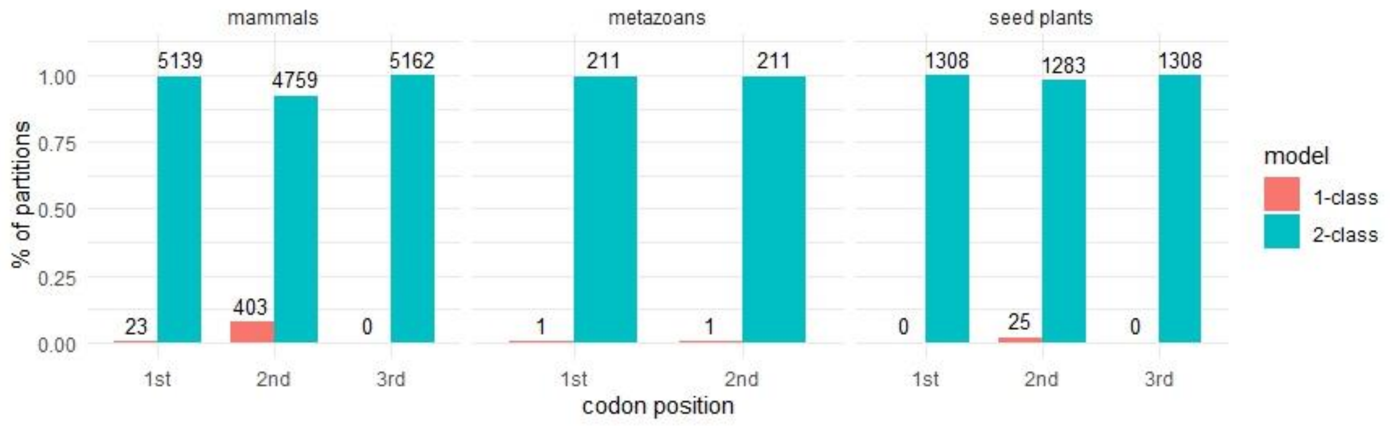

**Figure S3 Scatter plot of the normalized Robinson-Fould (nRF) distance between the commonly accepted tree and each of the 2-class tree and 1-class tree on each partition.** The red line (y = x) represents that the nRF distances of 1-class and 2-class trees to the concatenated tree are the same. The x axis is the nRF distance of 1-class tree to the concatenated tree and the y axis is the nRF distance of 2-class tree to the concatenated tree.

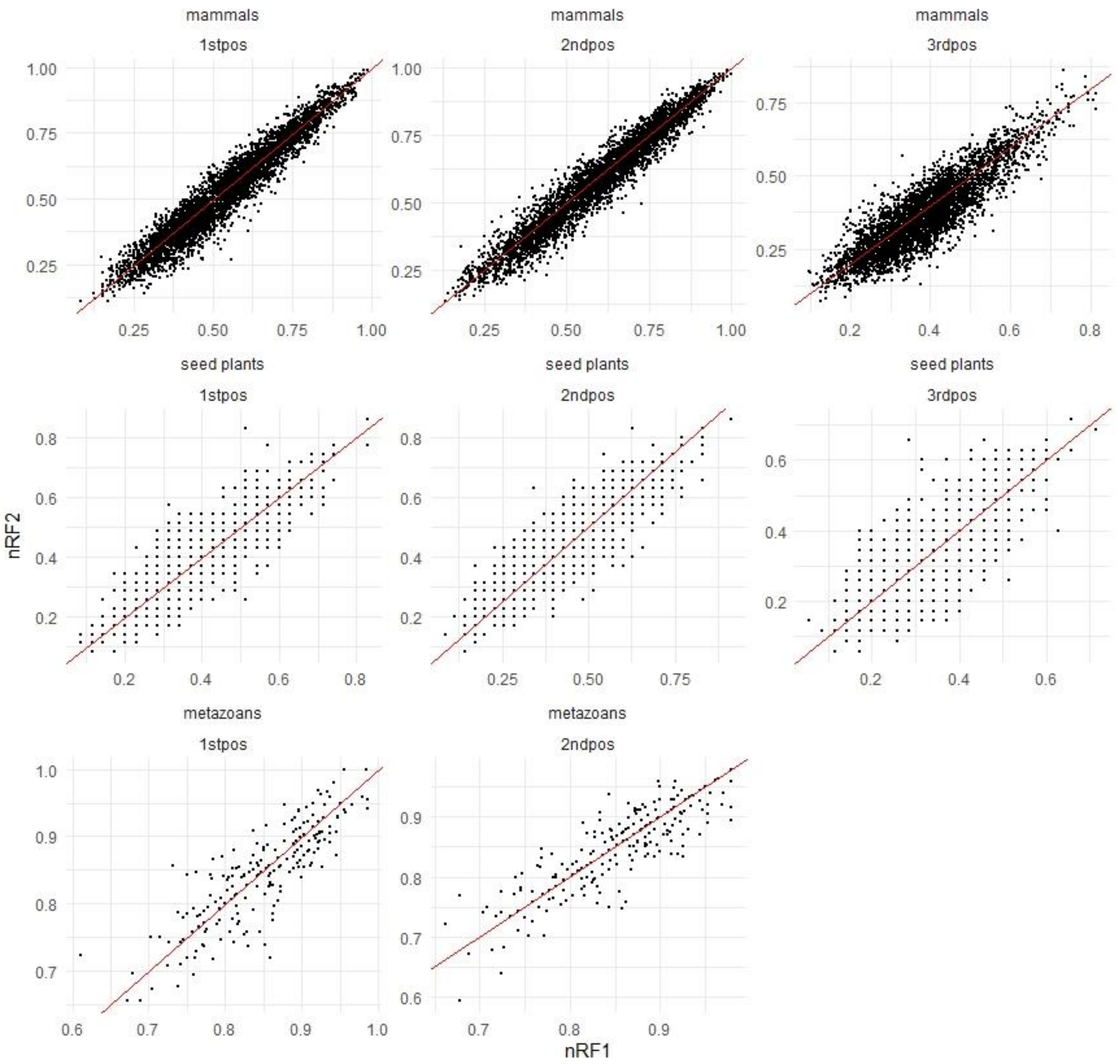

**Figure S4 Estimation of the number of model classes.** The red dashed line (y = x) represents that the estimated number of classes equals to the true number. The mean value and the corresponding standard error of the mean (error bar) of each experimental condition are shown with different curves.

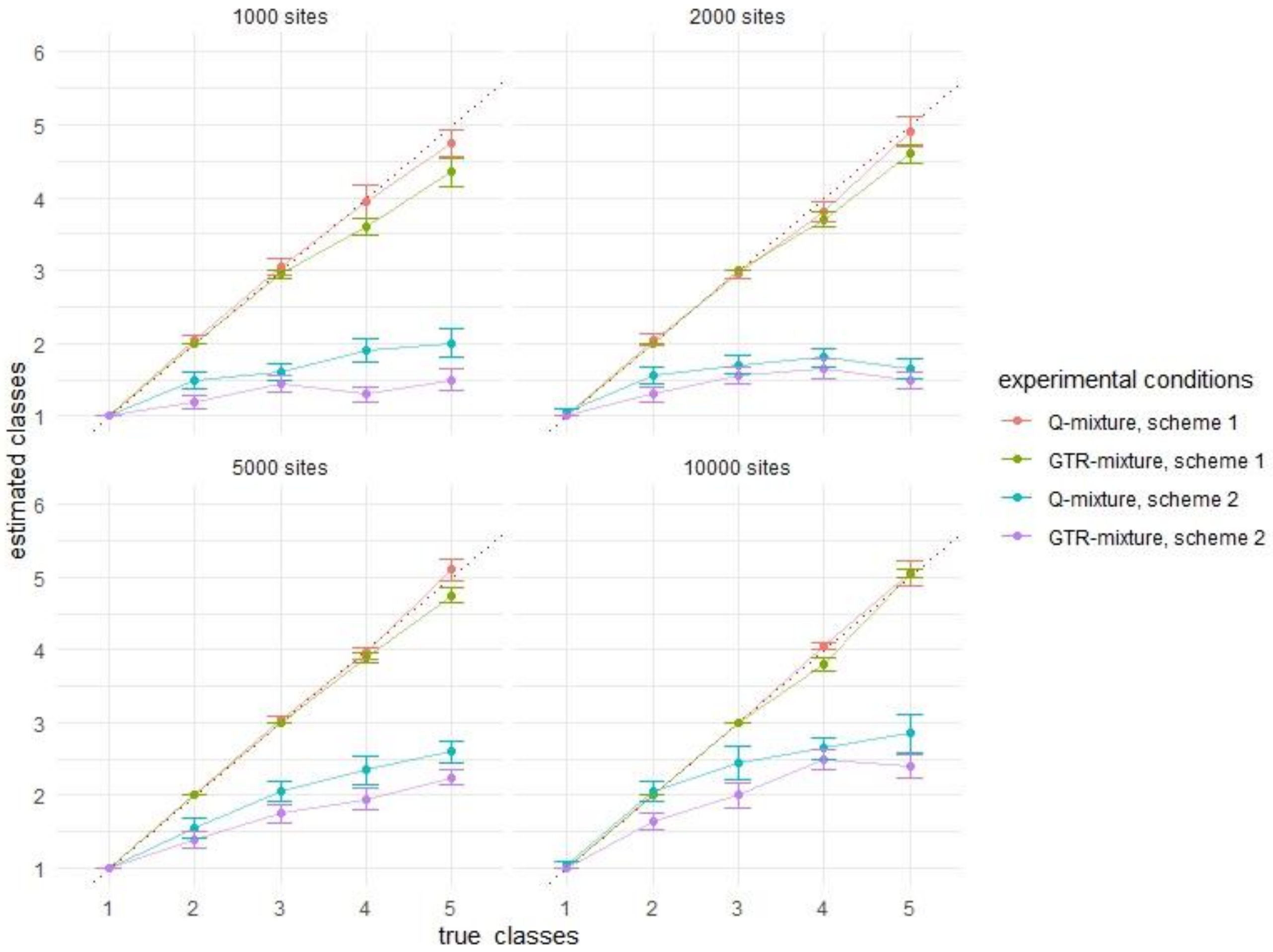

**Figure S5 Integrated Squared Error (ISE) between parameters in estimated model and simulated model.** The mixture models (green and blue bars) have smaller ISEs to the simulation than the non-mixture models (red bars). The “*” marks represent the significance of the paired t-test between Q-mixture and GTR-mixture models (0 '****' 0.0001 '***' 0.001 '**' 0.01 '*' 0.05).

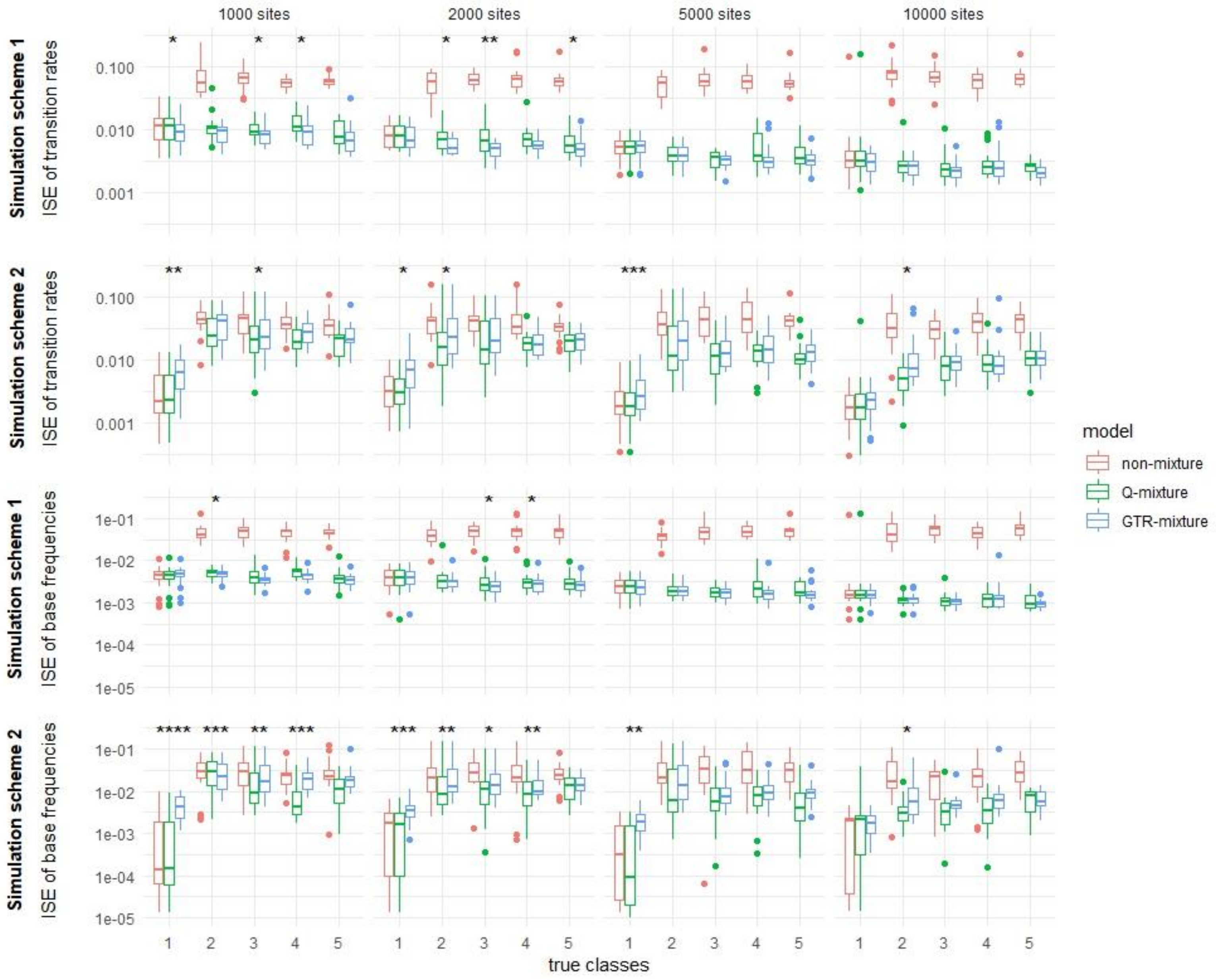

**Figure S6 The normalized Robinson-Fould (nRF) distance and relative difference in tree length between estimated models and simulated models.** The mean value and the corresponding standard error of the mean (error bar) of each experimental condition are shown with different curves. The nRF of mixture model trees to the simulated trees are slightly smaller than the non-mixture model in only simulation scheme 1, 10000-site alignments. In simulation schemes 1, the mixture models recover the correct tree lengths, while the non-mixture models underestimate the tree lengths. In simulation scheme 2, this tendency is only shown in 10000-site datasets.

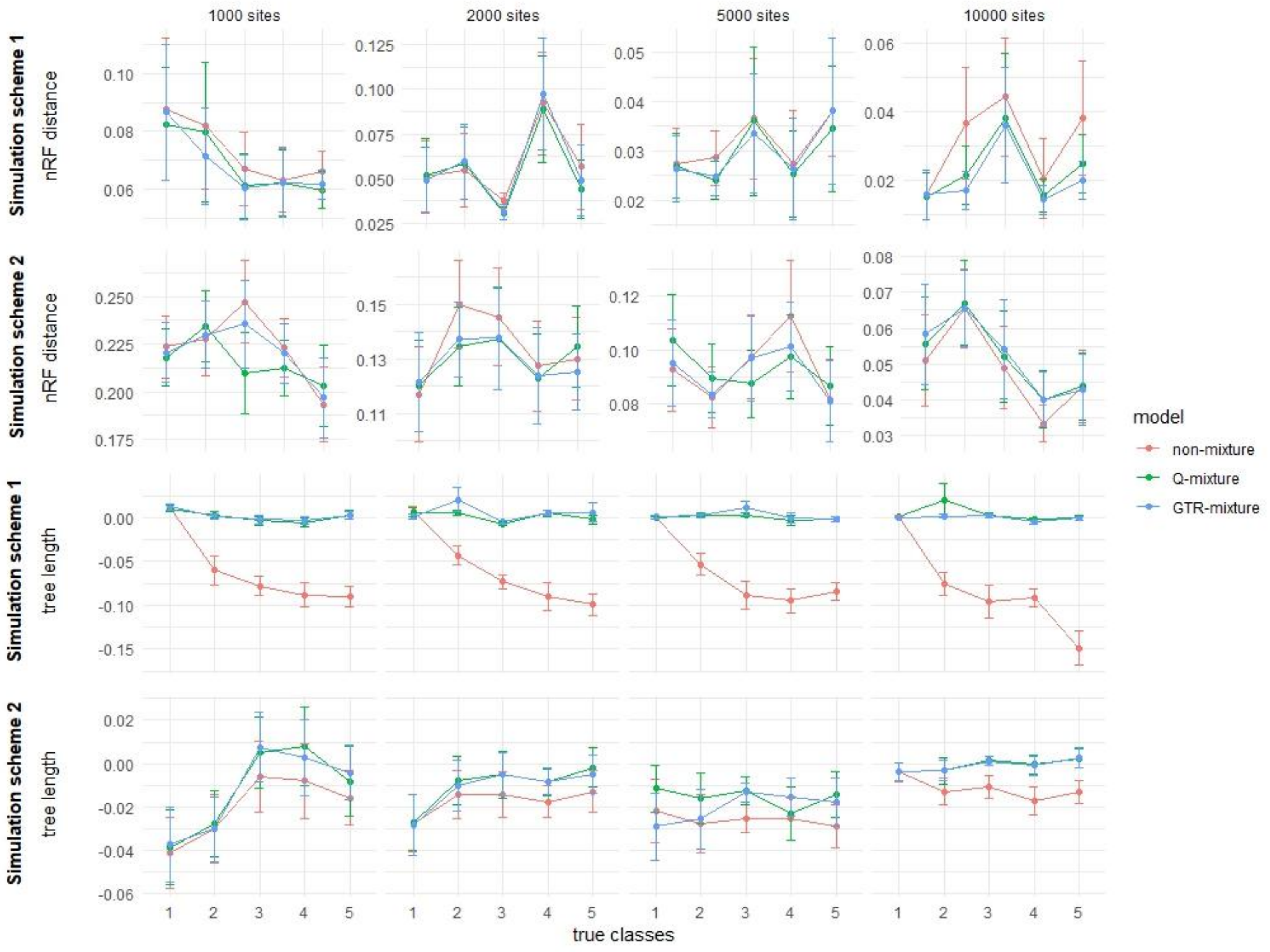

**Figure S7 Runtime and memory requirement of MixtureFinder.** We ran our simulation experiment with one thread for each dataset and measured the runtime and the maximum memory requirement of Q-mixture model and GTR-mixture model. Since the estimation of the number of classes was not always correct, the runtime and memory requirement do not always scale as expected with the number of simulated classes.

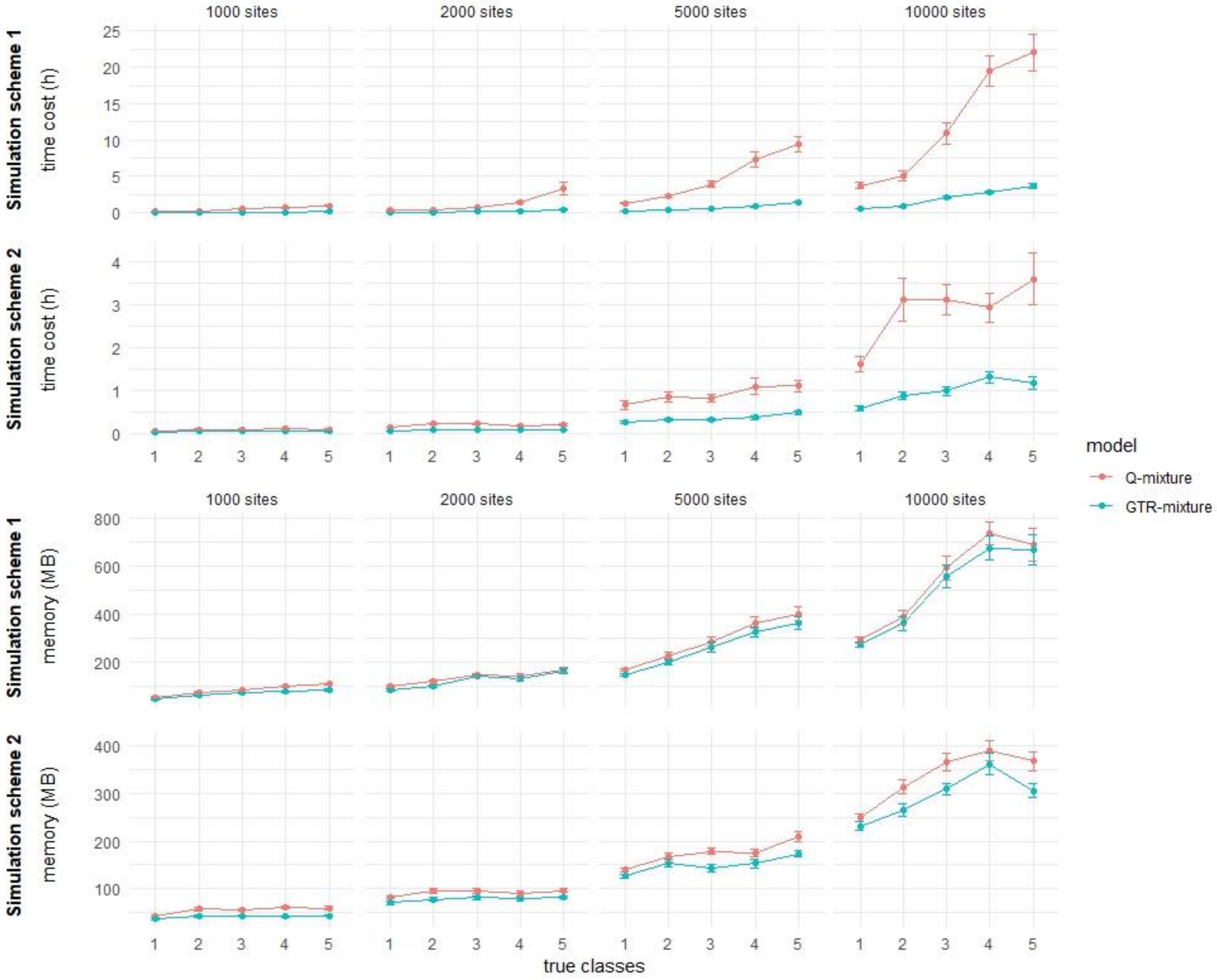

**Figure S8 Assessment of the class weight estimation on simulation.** We fit the mixture model with the correct number of classes with GTR model for each mixture class to each the simulated alignments described in Materials and methods of the main texts. This simulation contains two schemes: 1. Each Q matrix is a GTR model. 2. Each Q matrix are different and simulated based the empirical data. To align the classes of estimated mixture model and simulated true model, we calculate the root mean squared error (RMSE) between the parameters of each Q matrix in the estimated and simulated mixture model, then use Hungarian Algorithm(Munkres 1957) to find the alignment with the least total RMSE of the aligned Q matrix pairs in the mixture classes. Finally, we calculate the RMSE between the aligned estimated class weights and simulated class weights.

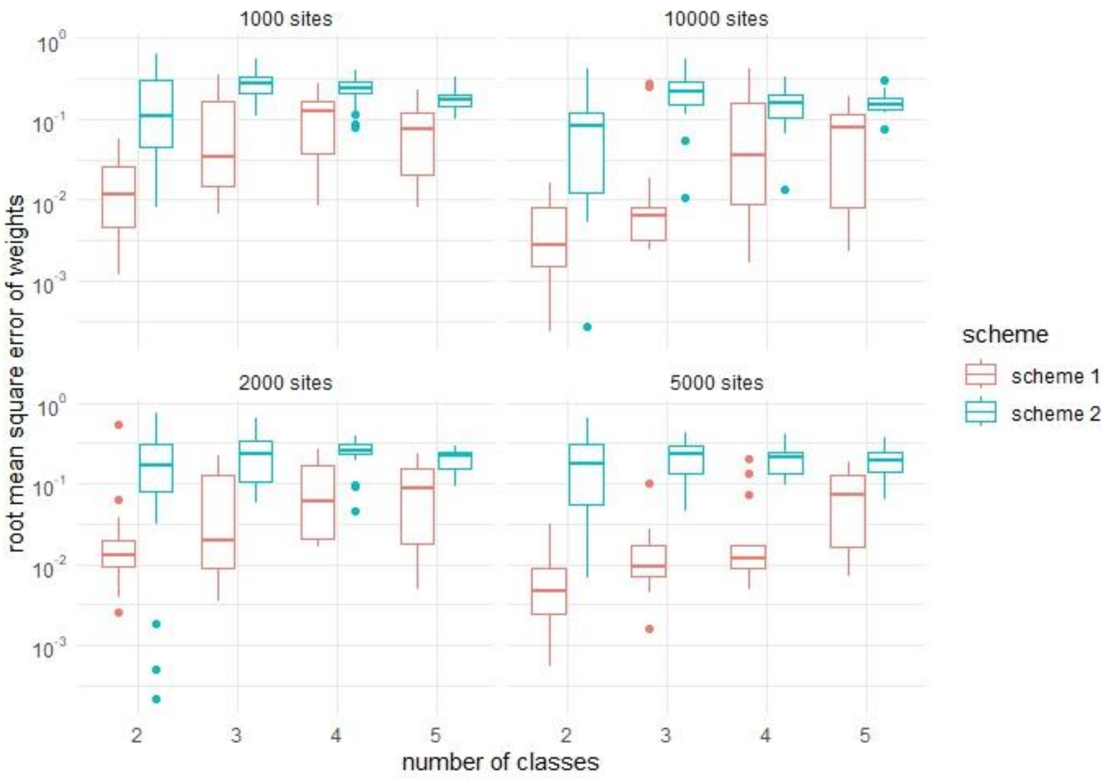

**Figure S9 Distribution of Likelihood Ratio Statistics (LRS) for comparing 2-class and 1-class models on 1-class simulation data.** The LRS values for 1000 dataset samples are shown in different facets based on different Q matrix in the second mixture class (JC, K2P, TPM3, HKY, TIM and GTR have 1, 2, 3, 5, 7 and 9 more degree of freedom respectively (including class weight)). **Red line:** the 95th percentile of the LRS. **Blue line:** the 0.05-level chi-squared test threshold with the degree of freedom that corresponding to each model.

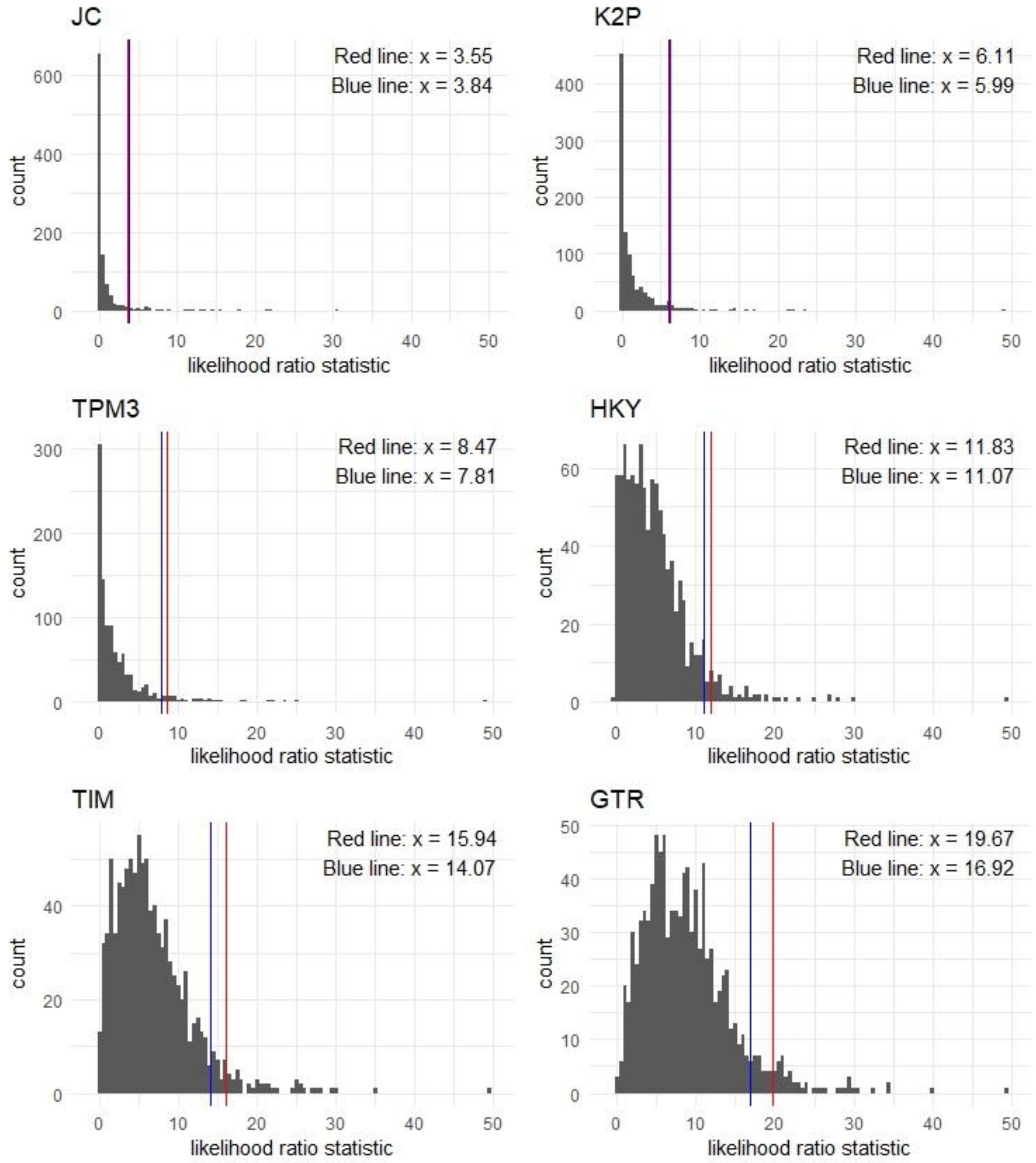

**Figure S10 Distribution of the Pythia-based difficulty score on the simulation data.** The density of the score based on different simulation schemes and different alignment lengths are shown respectively.

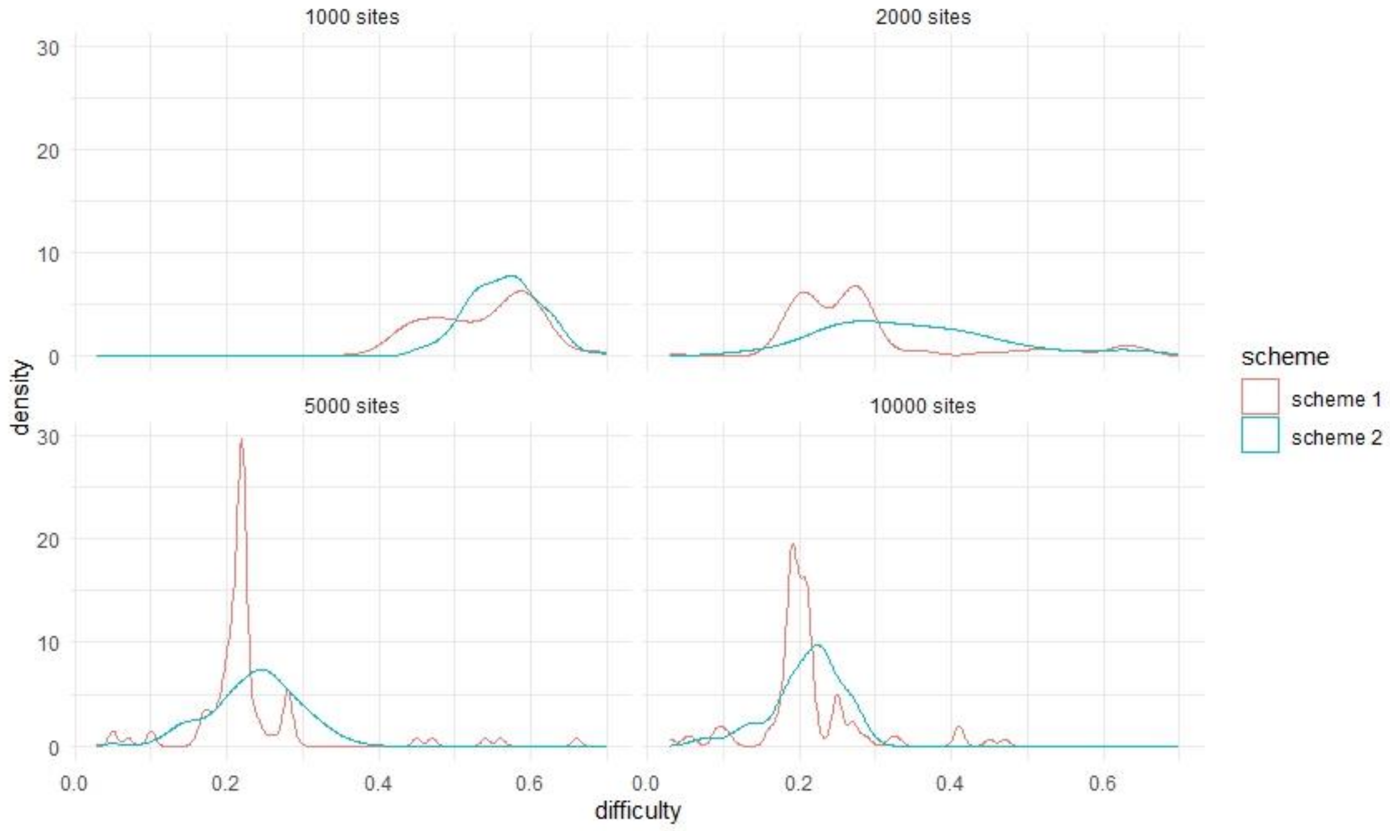

**Reference:**

Munkres, J. 1957. Algorithms for the assignment and transportation problems. *Journal of the Society for Industrial and Applied Mathematics* 5(1), pp. 32–38.
